# Abundance, diversity and evolution of tyrosinase enzymes involved in the adhesive systems of mussels and tubeworms

**DOI:** 10.1101/2024.07.05.602216

**Authors:** Emilie Duthoo, Jérôme Delroisse, Barbara Maldonado, Fabien Sinot, Cyril Mascolo, Ruddy Wattiez, Pascal Jean Lopez, Cécile Van de Weerdt, Matthew J. Harrington, Patrick Flammang

## Abstract

The blue mussel (*Mytilus edulis*) and the honeycomb tubeworm (*Sabellaria alveolata*) have evolved similar adhesive systems to cope with the hydrodynamic conditions of the intertidal environment where they live. Both organisms can establish a permanent adhesion through the secretion of adhesive proteins rich in DOPA (3,4-dihydroxyphenylalanine), a post-translationally modified amino acid playing essential roles in interfacial adhesion and bulk cohesion. DOPA is produced by the hydroxylation of tyrosine residues by tyrosinase enzymes, which can also in some cases oxidise it further into dopaquinone Compared to the detailed knowledge available on mussel and tubeworm adhesive proteins, little information exists about the tyrosinases involved in their adhesive systems. By combining different molecular analyses, a catalogue of tyrosinase candidates potentially involved in the adhesive systems of *M. edulis* and *S. alveolata* was identified. Some of these candidates were shown to be expressed in the adhesive glands by *in situ* hybridization, with a high gland-specificity in mussels but not in tubeworms. The diversity of tyrosinases highlighted in the two species suggests the coexistence of different functions (monophenol monooxygenase or catechol oxidase activity) or different substrate specificities. However, the exact role of the different enzymes needs to be further investigated. Phylogenetic analyses support the hypothesis of independent expansions and parallel evolution of tyrosinases involved in adhesive protein maturation in both lineages, supporting the convergent evolution of their DOPA-based adhesion.

## Introduction

Many marine organisms have evolved diverse attachment strategies to cope with their hydrodynamic environment (Delroisse *et al*. 2023). In particular, marine invertebrates rely on proteinaceous underwater adhesives for both permanent and temporary attachment to surfaces in the intertidal zone (Almeida *et al*., 2020; Li *et al*., 2021). These adhesives are known for their superior strength and durability compared with man-made materials and can therefore provide inspiration for the design and implementation of robust underwater adhesive strategies (Hofman *et al*., 2018). Two of the most extensively investigated organisms in this context are mussels and tubeworms (Fig. 1A-C). To attach themselves to rocks, mussels produce a byssus, which consists of a set of protein threads, each connected proximally to the base of the animal’s foot (Fig. 1B), inside the shell, and ending distally in a flattened plaque sticking to the substratum (Waite, 2017). Byssal threads are formed by the auto-assembly of proteins secreted by three distinct glands enclosed in the mussel foot: the plaque gland, the core gland, and the cuticle gland (Waite, 2017; Priemel *et al*., 2017). To build and expand the tube in which they live, tubeworms of the family Sabellariidae collect mineral particles from their surroundings, dab them with spots of cement, and then add them to the opening of their tube (Vovelle, 1965; Stewart *et al*., 2004). The cement consists mostly of several different proteins produced by two types of parathoracic unicellular glands (cells with homogeneous granules and cells with heterogeneous granules) (Fig. 1C) (Becker *et al*., 2012; Wang & Stewart, 2012). The presence of DOPA (3,4-dihydroxy-L-phenylalanine) is a distinctive feature common to proteins identified in the adhesive systems of both mussels and tubeworms (Stewart *et al*. 2011; Petrone 2013; Davey *et al*. 2021). This post-translationally modified amino acid fulfils crucial roles in both interfacial adhesive and bulk cohesive interactions within the adhesive secretions (Sagert *et al*. 2006; Waite, 2017). DOPA is produced by the post-translational hydroxylation of tyrosine residues of the adhesive proteins by tyrosinase enzymes (Silverman and Roberto, 2007; Priemel *et al*., 2020).

**Figure 1.**
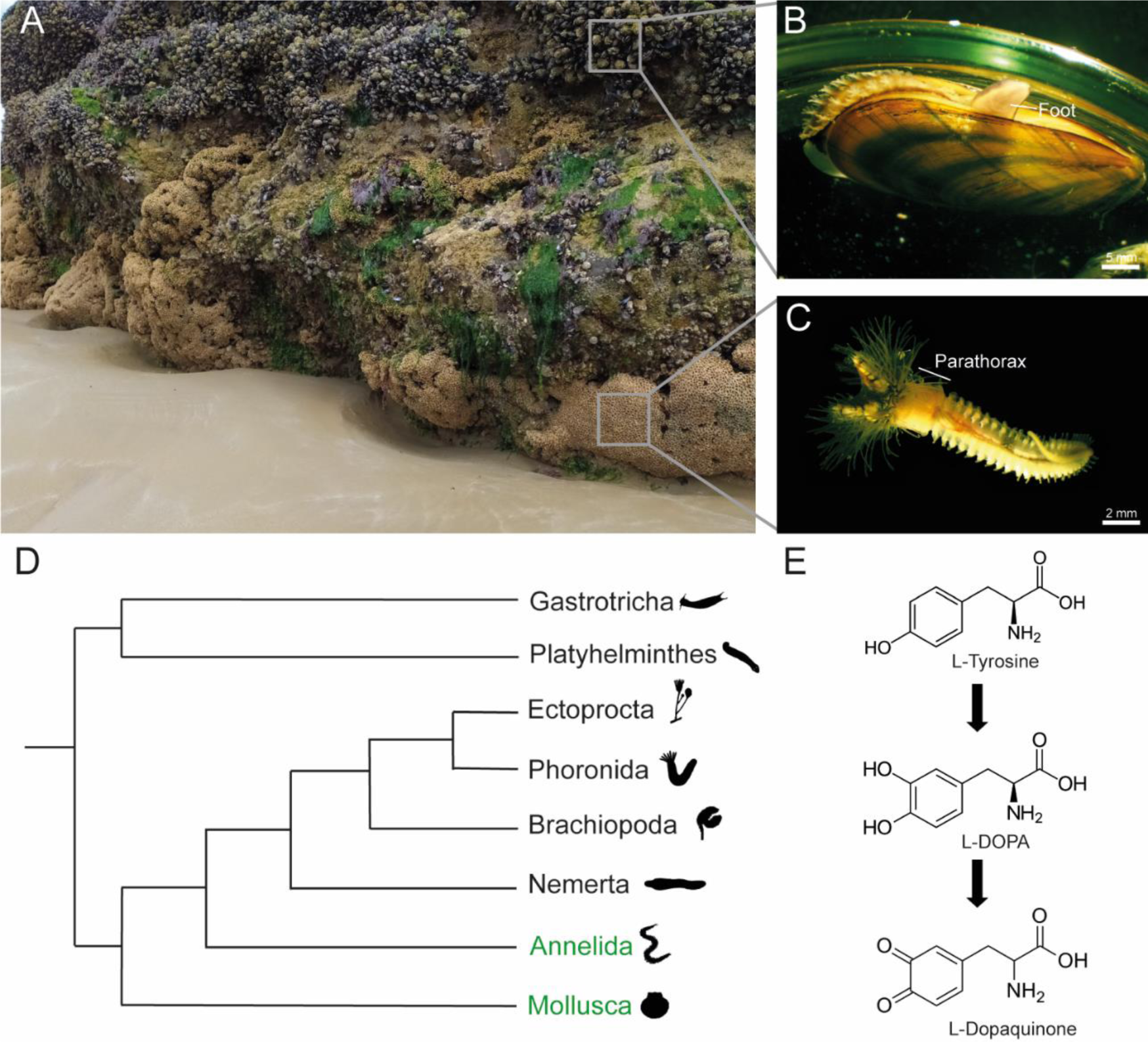
(**A**) Intertidal zone with a honeycomb worm reef covered by blue mussels (Douarnenez, France) (Picture courtesy of Alexia Lourtie). (**B, C**) *Mytilus edulis* and *Sabellaria alveolata*, respectively. (**D**) Phylogenetic position of mussels (Mollusca) and tubeworms (Annelida) (phyla written in green) within the Spiralia (based on Laumer *et al*., 2015). (E) The possible dual function of tyrosinases in the maturation of adhesive proteins comprising the modification of tyrosine residues into DOPA and the further oxidation of DOPA into dopaquinone (Polivares & Solano, 2009).

Tyrosinases belong to the type-3 copper protein family alongside hemocyanins (Aguilera *et al*., 2013). These oxygen-transferring copper metalloproteins catalyse the *o*-hydroxylation of monophenols (e.g., tyrosine) into o-diphenols (e.g., DOPA) and the further oxidation of *o*-diphenols to *o*-quinones (Fig. 1D) (Ullrich & Hofrichter, 2007; Aguilera *et al*., 2013). Consequently, these enzymes exhibit both cresolase (monophenol monooxygenase, EC 1.14.18.1) and catecholase (catechol oxidase, EC 1.10.3.1) activities (Ramsden and Riley, 2014). Tyrosinases are distributed throughout the tree of life (Aguilera et al., 2013). They are among the most widespread enzymes in nature as they are key enzymes for pigment synthesis in many organisms (Del Marmol & Beermann, 1996). They are also involved in many other biological processes such as shell formation in molluscs (Huan *et al*., 2013) or wound healing (Sugumaran *et al*., 1996; Theopold *et al*., 2004), parasite encapsulation (González-Santoyo & Córdoba-Aguilar, 2012) and cuticle sclerification (Dennell R., 1958; Andersen, 2010) in insects. Regardless of their biological function, all tyrosinases share a common origin traceable to an ancestral tyrosinase gene (Polivares & Solano, 2009). Lineage-specific gene duplication events took place during evolution, potentially contributing to functional diversification in specific taxa (Esposito *et al*., 2012; Aguilera *et al*., 2013).

Compared to the detailed knowledge available on mussel and tubeworm adhesive proteins, little information exists about the tyrosinases present in the adhesive systems of these organisms. In the mussel *Mytilus edulis*, Waite (1985), and later Hellio and collaborators (2000), reported the partial purification and kinetics of tyrosinases extracted from the foot and the byssus and displaying a catechol oxidase activity. More recently, developments in transcriptomics and proteomics allowed the discovery of new foot- or byssus-specific tyrosinase sequences. Guerette *et al*. (2013) identified 5 isoforms from the foot of the green mussel *Perna viridis* while Qin *et al*. (2016) detected 6 isoforms in the byssus of *Mytilus coruscus*. Their distribution within the foot or the byssus suggests they might have specific substrates and or functions. In tubeworms, one tyrosinase was identified in *Phragmatopoma californica* and shown to be expressed in the two types of adhesive glands (Wang & Stewart, 2012). A later study from the same authors demonstrated that the enzyme was a catechol oxidase that catalyses the covalent cross-linking of L-DOPA (Wang & Stewart, 2013). In two other species, *Phragmatopoma caudata* and *Sabellaria alveolata*, a differential transcriptomic study highlighted 23 tyrosinase transcripts overexpressed in the region of the body enclosing the adhesive glands (Buffet *et al*., 2018). Many studies therefore point to an expansion of the tyrosinase repertoire linked to the maturation of adhesive proteins in both mussels and tubeworms.

The adhesive secretions of mussels and tubeworms could represent model molecular systems for studying the evolution of tyrosinases and the diversification of their functions. Despite belonging to two distinct phyla, mussels and tubeworms are phylogenetically close, both being lophotrochozoans (Fig. 1E). Moreover, co-occurring in some intertidal habitats, they are subjected to the same environmental conditions and selective pressures, and their adhesive mechanisms are notably similar, primarily relying on DOPA. Several reviews have hinted at this similarity (Endrizzi & Stewart, 2009; Flammang *et al*., 2009; Stewart *et al*., 2011; Hofman *et al*., 2018). However, due to the short and intrinsically disordered nature of the adhesive protein sequences in both mussels and tubeworms, no homology can be traced between them, and they are thought to have evolved independently (Kamino, 2010). Examining the enzymes involved in the maturation process of adhesive proteins, especially tyrosinases, in these two taxa could provide insights into the evolutionary relationships between their adhesive systems. In this study, we investigated the catalogue of tyrosinases potentially involved in the maturation of adhesive proteins in the blue mussel *M. edulis* and the honeycomb worm *S. alveolata* by performing proteotranscriptomic analyses. *In situ* hybridization was then used to confirm the expression of these candidate enzymes in adhesive glands, validating their role in bioadhesion. Finally, we conducted phylogenetic analyses to address the question of the evolution and diversification of tyrosinases in these two lineages.

## Results

### Transcriptomic analyses

The initial phase of this study aimed at identifying enzymes containing a tyrosinase domain that might play a role in adhesive protein maturation in both *M. edulis* and *S. alveolata*. To achieve this, a mussel foot transcriptome and a honeycomb worm anterior part transcriptome were obtained. Both transcriptomes were generated from 100 bp reads sequenced using the Illumina platform. The raw data comprises 6.1 Gbp for *S. alveolata* and 3.9 Gbp for *M. edulis*. The *M. edulis* dataset contains 173,165 predicted genes, encompassing a total of 152,956,891 nucleotides. The contig N50 is 1,652 nucleotides, with a maximum contig length of 19,275 nucleotides. The *S. alveolata* dataset includes 414,599 predicted genes, amounting to 332,743,778 nucleotides in total. The contig N50 is 1,409 nucleotides, and the maximum contig length is 31,799 nucleotides. The sequence length distribution of the predicted transcripts is shown in the Supplementary Figure 1A-B. The completeness of both transcriptome datasets was evaluated using BUSCO (Benchmarking Universal Single-copy Orthologue) analyses on assembled transcripts. Scores were evaluated using the predefined lineage data “Metazoan_odb10”. For the *M. edulis*, analyses showed that 92.4% complete BUSCO groups were detected in the gene set. In detail, out of the 954 evaluated BUSCOs from the Metazoan dataset, only 5.2% were fragmented and 2.4% were missing. For *S. alveolata*, the BUSCO analyses indicated that 97.8% complete BUSCO groups were detected with only 2.2% and 0% of fragmented and missing BUSCO groups (Supplementary Figure 1C). Transcriptome data were then used to search for tyrosinase mRNA sequences using a similarity-based approach. A dataset of tyrosinases comprising mussel byssus (*Mytilus coruscus* [Qin *et al*., 2016], *Perna viridis* [Guerette *et al*., 2013]) and tubeworm cement (*Phragmatopoma californica* [Wang and Stewart, 2012]) sequences (see Suppl. Table 1) was used for local tBLASTn searches in the two transcriptomes to highlight sequences with similarity to these tyrosinases. Candidate matches were then used as queries in a reciprocal BLASTn search against online databases, and only those corresponding to a tyrosinase-like protein as the reciprocal hit were kept as putative candidates. Additionally, after *in silico* translation, short sequences and sequences lacking a tyrosinase domain (cl02830 or pfam00264) were removed.

In the blue mussel, the BLAST searches allowed retrieval of 85 transcripts coding for proteins with tyrosinases as the best reciprocal hit. This list was reduced to 17 candidates when only full-length or almost full-length sequences comprising a tyrosinase domain were considered, of which 8 are the closest homologues (blast best hits) of the reference tyrosinases from *M. coruscus* and *P. viridis* (Table 1). Most of the 17 sequences are full-length, except for Med-TYR2 (comp79852) and Med-TYR3 (comp74994) which lack the C-terminal region. All these proteins also possess a signal peptide indicating they are likely secreted at the level of the foot, some of them presumably by the byssus forming glands. Yet, the expression level of the tyrosinase encoding transcripts, expressed as Fragments Per Kilobase of transcript per Million mapped reads (FPKM), is on average one to two orders of magnitude lower than that of transcripts coding for mfp-1, mfp-2 and PreCol-NG, three byssal proteins produced by the cuticle, plaque and core glands, respectively (Table 1).

**Table 1.** List of tyrosinase sequences identified in the mussel *Mytilus edulis* after proteo-transcriptomic analyses (see text for details). Proteins with names in bold are the closest homologues to reference tyrosinases from *M. coruscus* and *P. viridis*. Indicated are the transcript ID form the foot transcriptome, the normalized expression level of the transcripts in the transcriptome (FPKM), the protein length in amino acid (with asterisks indicating incomplete sequences), the presence of a signal peptide, the peptide coverage of the translated transcripts from the MS-MS analysis (number of detected peptides in induced byssal threads [BT], distal foot tissues [DF] and proximal foot tissues [PF]), the position in the phylogenetic tree (Fig. 3), and top reciprocal blast. Three byssal proteins representative of the cuticle (Mfp-1), plaque (Mfp-2) and core (PreCol-NG) glands are included for comparison.

| Sequence name | Clade in phylogenetic tree | Transcript ID | FPKM | Lengt h (aa) | Signal peptide | Number of peptides identified by proteomic analyses |  |  | Reciprocal hit tBLASTn |  |
| --- | --- | --- | --- | --- | --- | --- | --- | --- | --- | --- |
|  |  |  |  |  |  | BT | DF | PF | Name | Accession number |
| Tyrosinases |  |  |  |  |  |  |  |  |  |  |
| Medu-TYR1 | I | comp83177_c0_seq2 | 510.1 | 702 | Y | 30 | 17 | 17 | *M. coruscus* byssal tyrosinase | KX268645 |
| Medu-TYR2 | I | comp79852_c1_seq2 | 59.2 | 678* | Y | 10 | 10 | 2 | *M. coruscus* byssal tyrosinase | KX268645 |
| Medu-TYR3 | I | comp74994_c0_seq1 | 8.1 | 489* | Y | 1 | / | / | *M. coruscus* byssal tyrosinase | KX268645 |
| Medu-TYR4 | I | comp80529_c1_seq1 | 32.9 | 686 | Y | / | / | / | *M. coruscus* byssal tyrosinase | KX268645 |
| Medu-TYR5 | I | comp87104_c0_seq1<br>1 | 64.7 | 805 | Y | / | / | 3 | *Mizuhopecten yessoensis* tyrosinase | XM_021518068 |
| Medu-TYR6 | I | comp75651_c0_seq1 | 17.83 | 802 | Y | / | / | / | *Mytilus californianus* uncharacterized LOC127724782 | XM_052231853 |
| Medu-TYR7 | II | comp83597_c2_seq1 | 12.6 | 580 | Y | / | / | / | *M. californianus* uncharacterized LOC127706830 | XM_052211537 |
| Medu-TYR8 | III | comp66244_c0_seq1 | 15.0 | 573 | Y | / | / | / | *M. californianus* tyrosinase-like | XM_052233075 |
| Medu-TYR9 | III | comp83819_c0_seq1 | 34.9 | 628 | Y | / | / | / | *M. californianus* uncharacterized LOC127706823 | XM_052211524 |
| Medu-TYR10 | IV | comp85123_c3_seq1 | 215.0 | 696 | Y | 26 | 3 | 12 | *M. coruscus* byssal tyrosinase | KP322726 |
| Medu-TYR11 | IV | comp76132_c0_seq1 | 80.9 | 648 | Y | 21 | 2 | 8 | *M. coruscus* byssal tyrosinase | KP322726 |
| Medu-TYR12 | IV | comp83122_c1_seq1 | 29.9 | 694 | Y | 20 | 16 | / | *M. coruscus* tyrosinase | KP876481 |
| Medu-TYR13 | IV | comp72405_c0_seq1 | 12.1 | 439 | Y | 1 | / | / | *M. coruscus* tyrosinase-like | KP757802 |
| Medu-TYR14 | IV | comp76510_c0_seq1 | 1.3 | 393 | Y | / | / | / | *M. californianus* tyrosinase-like | XM_052215114 |
| Medu-TYR15 | IV | comp79670_c1_seq2 | 192.8 | 561 | Y | / | / | / | *M. galloprovincialis* catechol oxidase | MG975894 |
| Medu-TYR16 | IV | comp83799_c1_seq1 | 66.0 | 490 | Y | / | / | / | *M. californianus* tyrosinase-like | XM_052240476 |
| Medu-TYR17 | IV | comp84313_c0_seq2 | 5.0 | 597 | Y | / | / | / | *M. californianus* tyrosinase-like | XM_052207170 |
| Other representative byssal proteins |  |  |  |  |  |  |  |  |  |  |
| Mfp-1 |  | comp48040_c0_seq1 | 2213.6 | 100* | N | / | / | / | *M. edulis* gene for polyphenolic adhesive protein | X54422 |
|  |  | comp63881_c0_seq2 | 0.38 | 165* | N | / | 2 | / | *Mytilus edulis* clone 21 foot protein 1 (fp-1) mRNA, complete cds | AY845258 |
| Mfp-2 |  | comp74249_c0_seq2 | 4210.2 | 508 | Y | 17 | 19 | / | *M. edulis* clone 7 foot protein 2 (fp-2) mRNA | AY845261 |
| PreCol-NG |  | comp75832_c0_seq1 | 1594,4<br>5 | 339 | Y | 12 | / | 5 | *M. edulis* nongradient byssal precursor, mRNA | AF414454 |

In the honeycomb worm, a total of 86 transcripts encoding tyrosinase-like enzymes were identified, among which 28 were (almost) full-length and, once translated *in silico*, comprised a tyrosinase domain. For *S. alveolata*, an additional filtering step was implemented based on the differential expression data reported in Buffet *et al*. (2018). Only transcripts overexpressed in the parathoracic region of the worms were considered, bringing the number of candidates down to 13 (Table 2). In this species too, most of the sequences were full-length, except for Salv-TYR1 (comp274293) which is incomplete in N-term and therefore lacks a signal peptide. Although the sequence of Salv-TYR3, the closest homologue to the catechol oxidase from *P. californica*, appeared to be complete in our transcriptome, no signal peptide could be detected. Similar to mussels, the expression level of mRNAs encoding tyrosinases is much lower than that of mRNAs encoding the cement proteins Sa-1, Sa-2, and Sa-3A/B (Table 2).

**Table 2.**
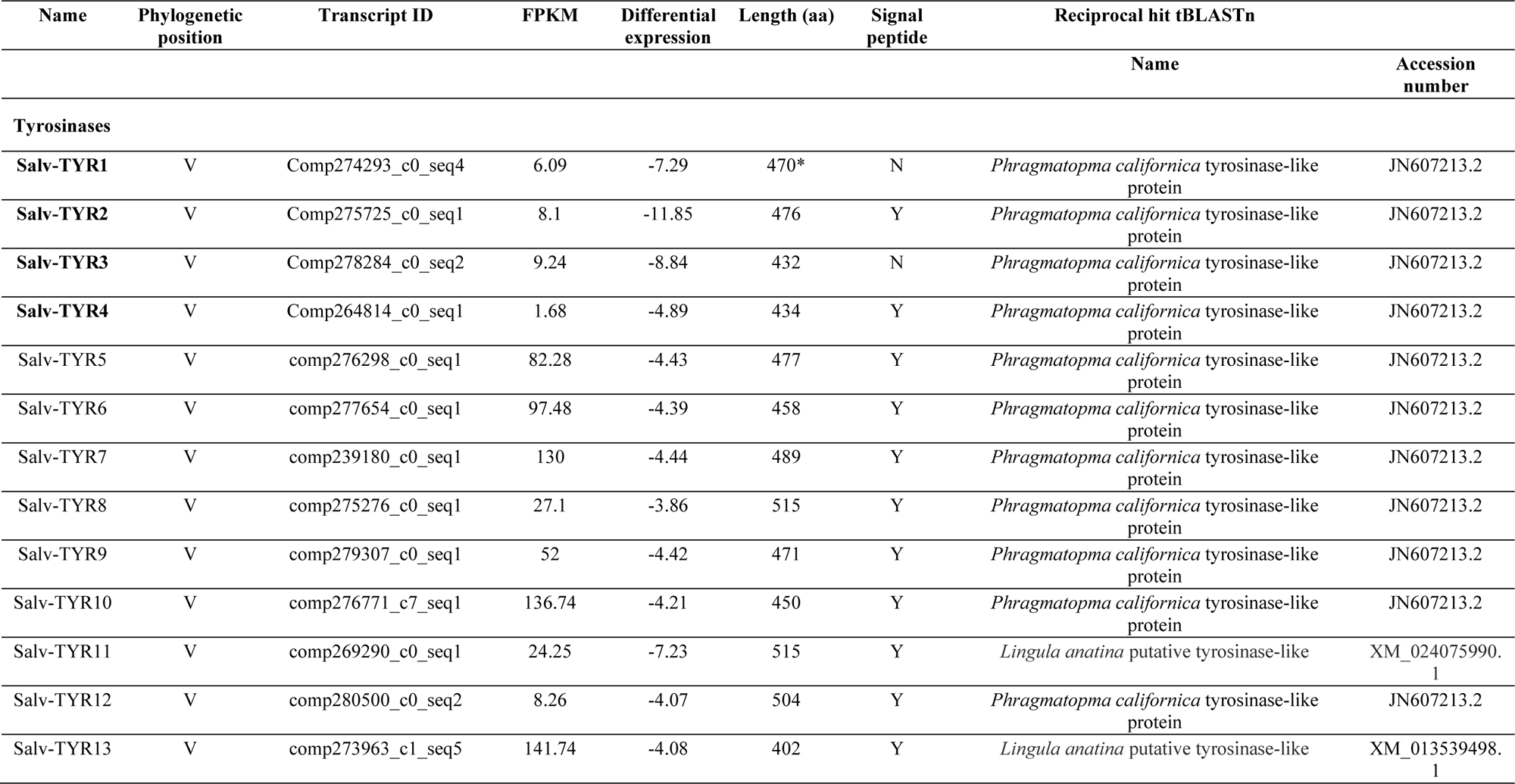
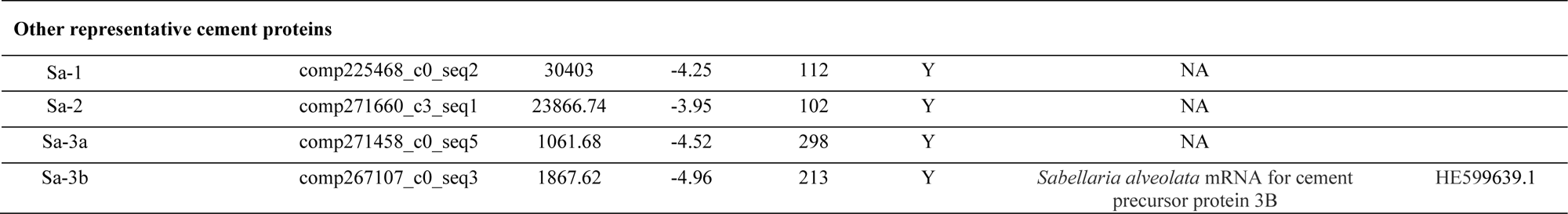
List of tyrosinase sequences identified in the tubeworm *Sabelleria alveolata* after transcriptomic analyses (see text for details). The protein with the name in bold is the closest homologue to the reference tyrosinase from *P. californica*. Indicated are the transcript ID from the transcriptome of the anterior part of the worm, the normalized expression level of the transcript in the transcriptome (FPKM), the differential expression of the transcript between the parathoracic part of the worm and the rest of its body (log2FoldChange reported in Buffet *et al*., 2018), the protein length in amino acid (with asterisks indicating incomplete sequences), the presence of a signal peptide, the position in the phylogenetic tree (Fig. 3), and the top reciprocal BLAST hit. Four cement proteins are included for comparison.

### Proteomic analyses

We carried out protein analyses on different samples from *M. edulis.* The secretion of byssal threads was induced by injecting KCl at the base of the foot. These freshly secreted threads were collected with fine forceps and the proteins comprising them were extracted with an 8M urea solution in 5% acetic acid and analysed by de novo peptide sequencing in mass spectrometry (MS/MS). In addition, mussel feet were dissected and cut in half to separate the foot tip from the basal part. Foot proteins were extracted with 4% sodium dodecyl sulfate (SDS) and were also analysed by MS/MS after trypsin digestion. By comparing the mass spectrometry results with the foot transcriptome, a total of 7 tyrosinases were identified with at least two peptides, with five detected in both the induced threads and foot samples, with a high peptide coverage, and two detected exclusively in the foot samples, but with only 3 peptides each (Table 1). Medu-TYR1, for example, exhibited the highest peptide coverage and its corresponding transcript was also the most abundant in the transcriptome compared to other candidates (Table 1). However, the correlation between abundance at the transcript and protein levels does not hold true for the other candidates. Among the tyrosinases identified in foot tissues, 2 were expressed exclusively in the tip (Medu-TYR8 and 12), 2 in the basal part (Medu-TYR5 and 10), and 3 in both parts (Medu-TYR1, 2 and 11). As for the byssal proteins used for comparison, two were detected (mfp-2 and preCol-NG) in all samples and their distribution in the foot corresponded to their expected expression pattern, only in the tip for mfp-2 and in both parts of the foot for preCol-NG (Table 1). Due to its tandemly repeated sequence making assembly difficult, Mfp-1 was represented in our transcriptome with two partial transcripts. Only two peptides corresponding to the sequence encoded by one of these transcripts could be detected by mass spectrometry.

In *S. alveolata*, the proteomic analysis was performed on tube fragments constructed by the worms using glass beads. Indeed, in the laboratory, isolated individuals can rebuild their tubes with different materials, including clean glass beads. Freshly constructed tube fragments were collected, and proteins were extracted from the cement dots present on the surface of glass beads using 7 M guanidine hydrochloride. Despite these highly denaturing conditions, no protein was detected in mass spectrometry, suggesting that the adhesive secretion might be highly crosslinked and therefore posed challenges for protein extraction.

### Phylogenetic analyses

To gain a comprehensive understanding of the relationships between tyrosinases, we investigated protein sequences containing the tyrosinase domain (pfam00264) from various spiralian species. These sequences were retrieved from the NCBI database. They include 30 proteins from Annelida, 8 from Brachiopoda, 375 from Mollusca, and 63 from Platyhelminthes. Additionally, we incorporated sequences obtained through the proteo-transcriptomic analyses of our two model species, specifically 20 predicted proteins from *M. edulis* (3 sequences found in the mantle were added to the 17 previously identified) and 28 from *S. alveolata* (Tables 1 and 2) (Suppl. Table 1). A sequence similarity-based clustering analysis was carried out using CLANS (Frickey & Lupas, 2004) to explore potential similarities between all these tyrosinases. An all-against-all BLASTp was conducted using the scoring matrix BLOSUM62 and linkage clustering was performed with a maximum E-value of 1E^−20^ to identify coherent clusters. The clustering was initially performed in 3 dimensions and then projected into 2 dimensions to generate the illustration shown in Fig. 2. The darker connections between the dots indicate higher similarity between the proteins based on the BLASTp E-values (Pearson, 2013).

**Figure 2.**
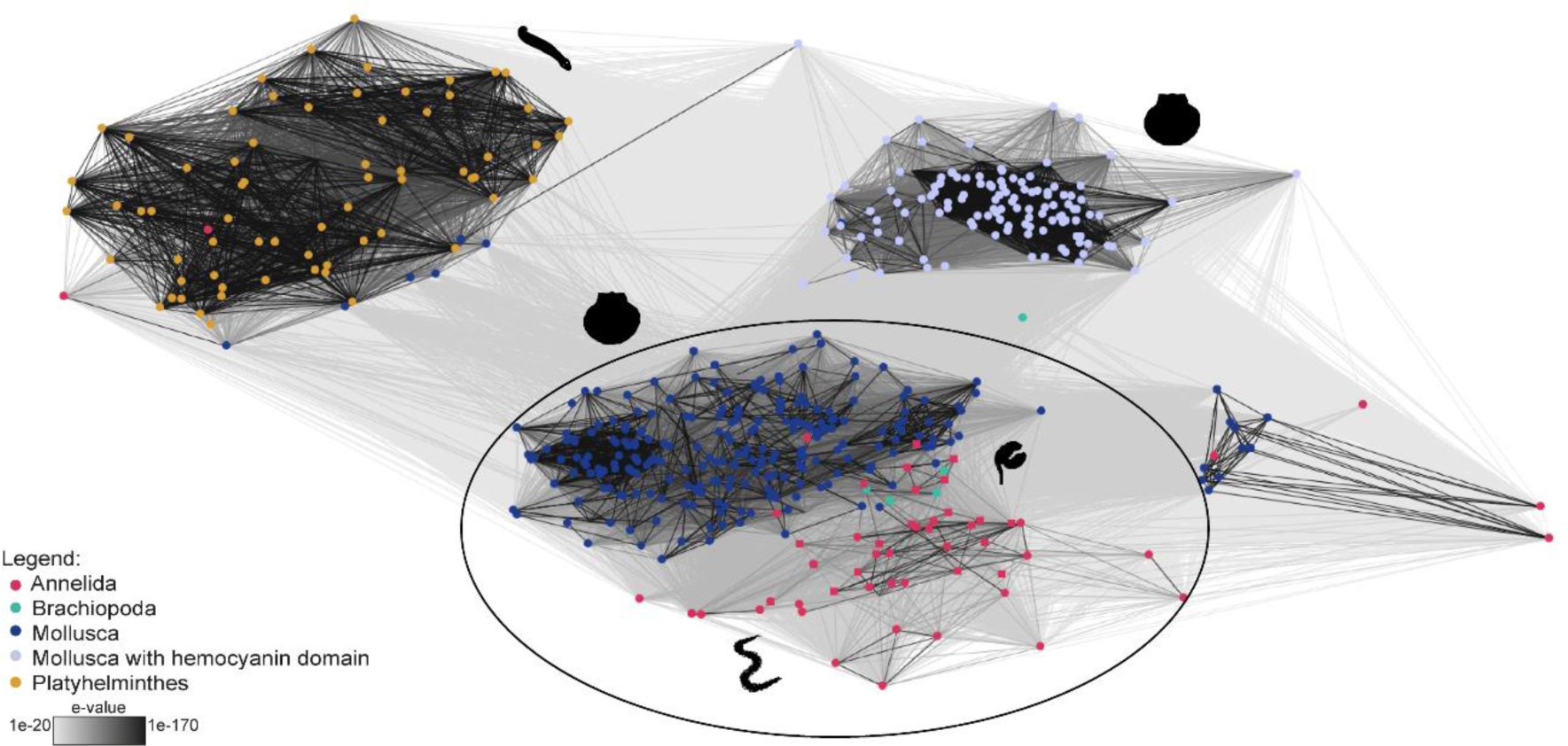
Sequence-similarity-based clustering approach based on BLASTp e-values with all the spiralian sequences containing a pfam00264 domain retrieved from the NCBI database. This analysis shows that the different phyla tend to cluster together. The sequences circled were selected for phylogenetic analyses.

The CLANS analysis revealed that spiralian tyrosinases form several clearly distinct clusters, each comprising sequences that share a high similarity among them as evidenced by the low E-values associated to the lines connecting them (Fig. 2). All the tyrosinase sequences from each phylum are generally grouped together, although a few isolated dots are noticeable, usually corresponding to very short (partial) sequences or very long sequences (>3000 amino acids) comprising additional domains (e.g., kielin/chordin-like domain). The phylum Mollusca is an exception as its sequences were divided into two clusters: one comprising sequences with only a tyrosinase domain and one consisting of sequences containing both a tyrosinase (pfam00264) and an hemocyanin domain (pfam14830). Hemocyanins are extracellular proteins involved in oxygen transport and are found in the phyla Arthropoda and Mollusca (Van Holde *et al*., 1995; Solomon *et al*., 1996). Previous studies have demonstrated that both hemocyanins and tyrosinases belong to the type 3 copper protein family and share a common ancestor (Drexel *et al*., 1987; Burmester & Schellen, 1996). However, structural modifications at the binuclear copper active site underlie the divergent evolution of tyrosinase and hemocyanin functions (Aguilera *et al*., 2013). This functional divergence may explain why all the Mollusca sequences are not grouped in the same cluster.

In the CLANS analysis, tyrosinase sequences from Platyhelminthes form a well-separated cluster whereas those from Lophotrochozoa (Brachiopoda, Annelida and Mollusca) are grouped together (Fig. 2). To delve deeper into their evolutionary relationships, the 312 tyrosinase protein sequences from the lophotrochozoan super-cluster were subjected to a phylogenetic analysis using molluscan hemocyanins as outgroup (Fig. 3, Suppl. Fig. 2). On the phylogenetic tree generated, 5 main clades (numbered from I to V on Fig. 3) can be distinguished. Clades I to IV group together the sequences from molluscs: clades I, II and IV contain only tyrosinases from bivalves, while clade III includes sequences from all classes of molluscs, from gastropods to cephalopods. Clade V contains a few oyster tyrosinase sequences (Bivalvia, Ostreoida), but comprise mostly non-molluscan sequences. Most of the sequences from the phylum Annelida are grouped with those from Brachiopoda in this clade, except some of the sequences from the polychaete *Owenia fusiformis* (Delle Chiaje, 1844) which are in clade III. The reference byssal tyrosinase sequences from the mytilids *M. coruscus* and *P. viridis*, as well as their closest homologues from *M. edulis* (Table 1) are localized either in cluster I (e.g., Medu-TYR1 to 4; Fig. 3) or in cluster IV (Medu-TYR10 to 15; Fig. 3). However, a few of the sequences shortlisted in the blue mussel are also present in clades II and III (Table 1). The reference cement tyrosinase from *P. californica* and all the sequences from *S. alveolata* are in cluster V. It should be noted that the phylogenetic tree generated from these analyses showed low support for some nodes (Fig. 3), likely due to the high level of conservation of residues surrounding the copper-binding sites and globally the short protein-based alignment. Similar results have been observed in previous studies (Aguilera *et al*., 2014; Buffet *et al*., 2018).

**Figure 3.**
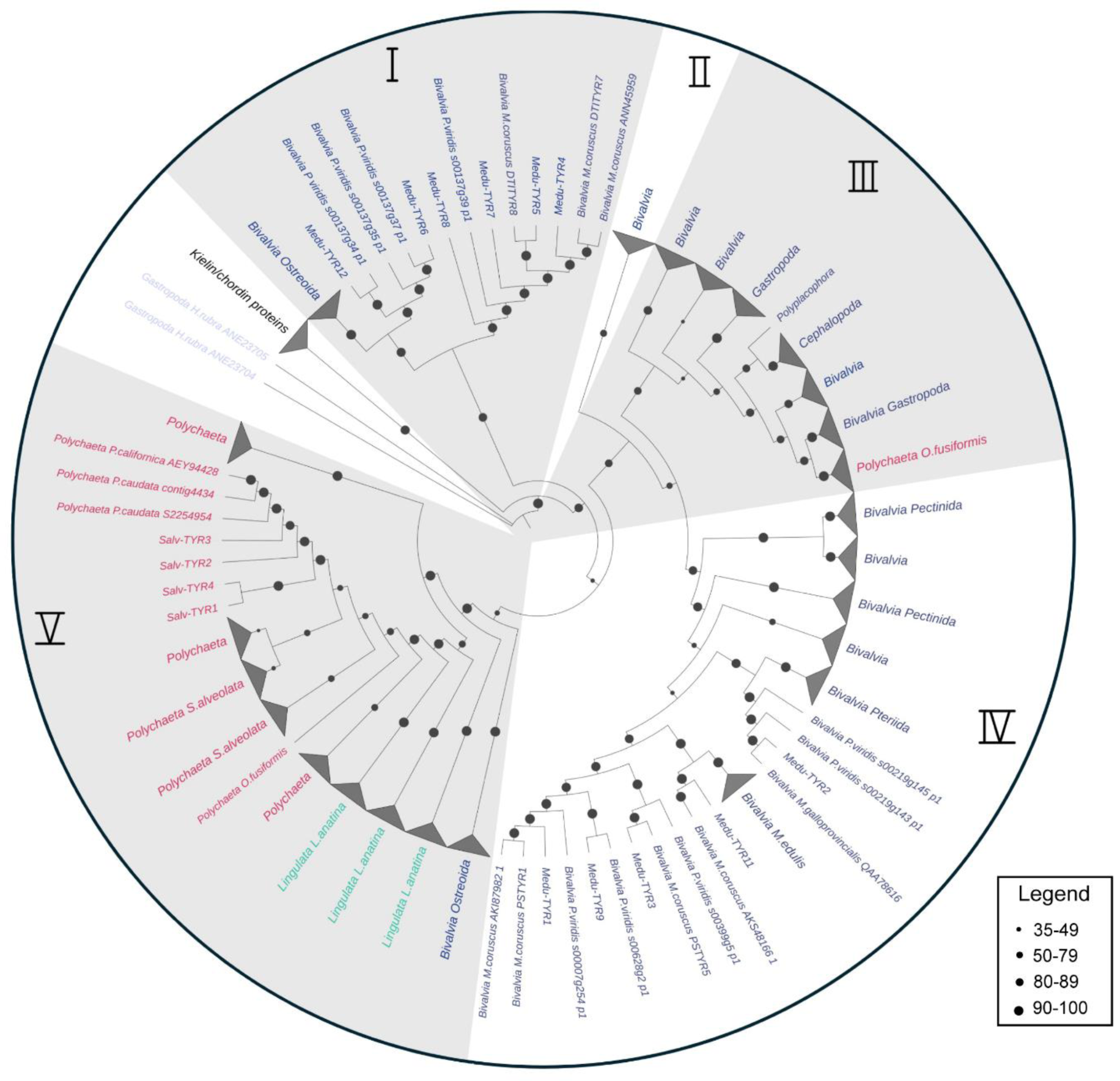
Maximum likelihood phylogenetic tree showing the distribution of tyrosinase sequences in Lophotrochozoa based on circled sequences selected in the CLANS analysis. The sequences are distributed among 5 main clades, numbered I to V. The reference byssal tyrosinase sequences from *M. coruscus* and *P. viridis* and their orthologues in *M. edulis* (Table 1) are localized in clusters I and IV, while the reference cement tyrosinase from *P. californica* and all sequences from *S. alveolata* (Table 2) are in cluster V. The bootstrap values are shown in the legend in the box.

### *In situ* hybridization experiments

To further investigate the role of the identified enzymes in adhesive protein maturation, the localization of their coding mRNAs in the tissues was determined using *in situ* hybridization (ISH). DIG-labelled RNA probes were designed based on sequences retrieved from the transcriptomes and used on tissue sections from both mussels and honeycomb worms, as well as on whole-mount preparations of the mussel’s feet, to determine whether the tyrosinases are expressed in the adhesive-producing gland cells or in other cell types. Controls were performed using sense RNA probes, as well as without probes or without antibody (Suppl. Fig. 3).

In the case of the blue mussel, we selected tyrosinase sequences showing the highest similarity to the reference sequences and/or coding for proteins detected with a minimum of two peptides in the induced byssal threads by MS/MS analyses. This corresponds to nine candidates: Medu-TYR1 to 5, Medu-TYR10 to 12, and Medu-TYR15 (Table 1). *In situ* hybridization was performed on the entire foot cut open in two halves along the central frontal plane (whole mount; see Fig. 4) but also on several transverse sections through the foot to make sure to visualize all specific foot glands, namely the core, cuticle, and plaque glands. For each section processed for ISH, a directly consecutive section was stained with Heidenhain’s azan to validate the identification of the glands. The results of the ISH experiments are illustrated in Fig. 4 and Suppl. Fig. 4. Among the nine candidates, one was found to be exclusively localised in the plaque gland (Med-TYR12), one in the core gland (Med-TYR10), and one in the cuticle gland (Med-TYR11). On the other hand, five candidates were expressed in at least two glands: both the plaque and the cuticle glands (Med-TYR4 and 15), both the plaque and core glands (Med-TYR2 and 5), or both core and cuticle glands (MeduTYR1). No probe could be produced for the candidate Med-TYR3.

**Figure 4.**
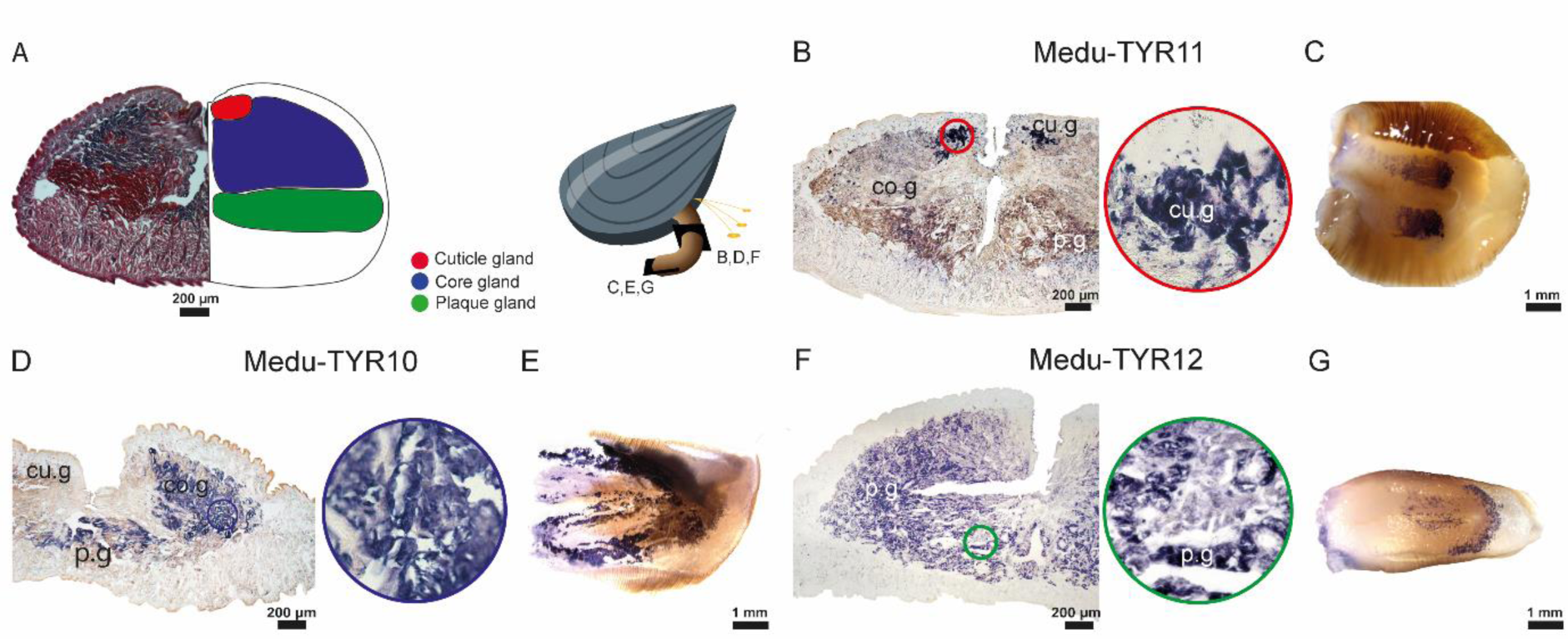
Examples of the expression patterns of selected tyrosinase transcripts in the foot of *Mytilus edulis* (**A**) Illustration showing both a transverse histological section of a foot stained with Heidenhain’s azan stain (left), and a diagram with the arrangement of the different glands in the foot tissues (right). (**B-G**) Section (B,D,F) and whole mount (C,E,G) *in situ* hybridization of three tyrosinase candidates. Insets in circles show a zoom on the labelled gland. Abbreviations: cu.g – cuticle gland; co.g – core gland; p.g – plaque gland.

For the honeycomb worm, four transcripts were selected: those exhibiting the highest similarity to the reference sequence from *P. californica* (Fig. 3), among which three were also the most highly differentially expressed in the parathoracic region of the worm (Table 2). The four tyrosinase candidates (Salv-TYR1 to 4) were exclusively expressed in the cement glands (Fig. 5). Moreover, they were all expressed within both cells with heterogeneous granules and cells with homogeneous granules (Fig. 5). The two types of granules can easily be distinguished at high magnification.

**Figure 5.**
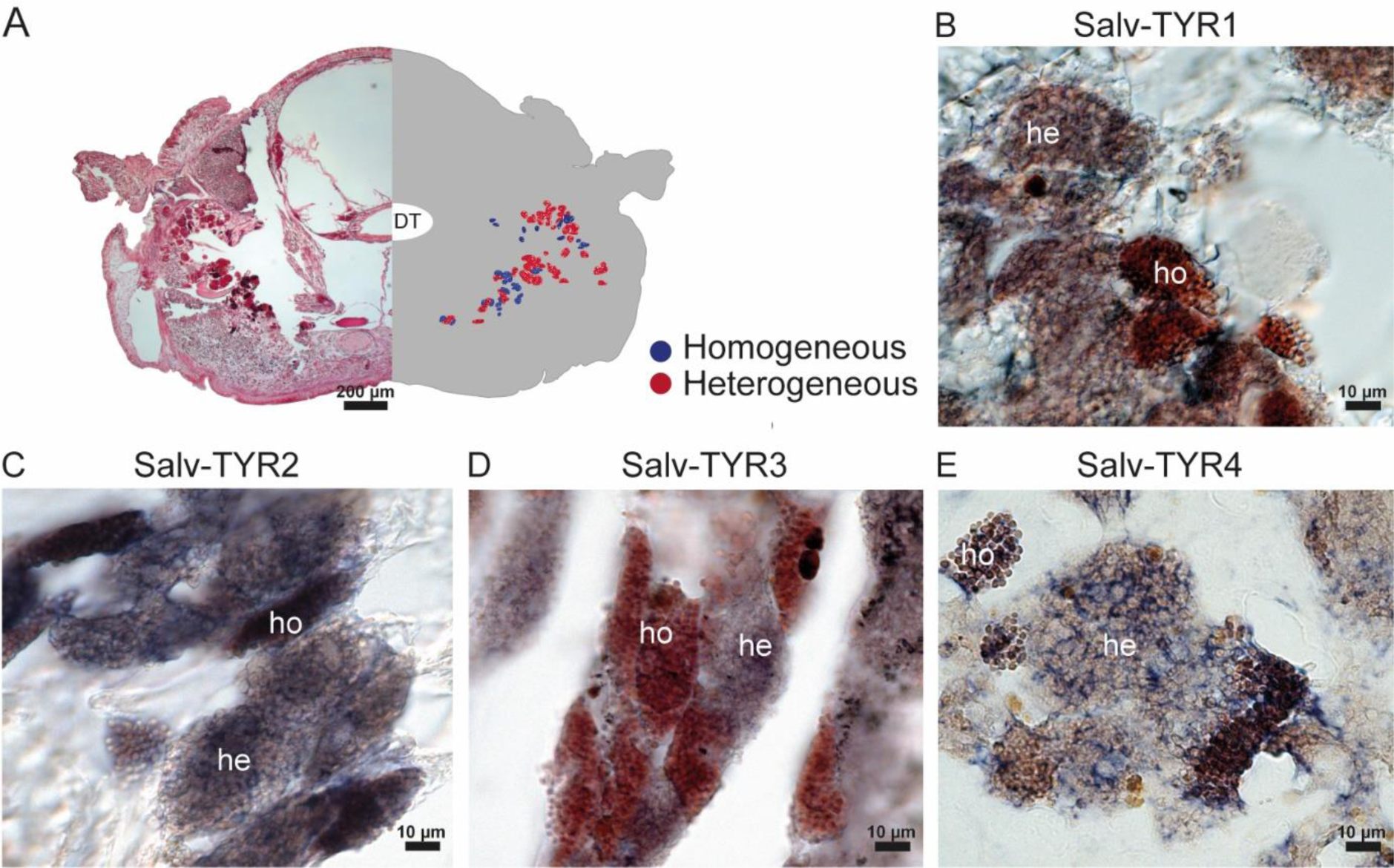
Expression patterns of selected tyrosinase transcripts in the parathoracic region of *Sabellaria alveolata*. (A) Illustration showing both a transverse section of the parathoracic part stained with Heidenhain’s azan stain (left), and a diagram showing the arrangement of the cement glands (right). (B-D) *In situ* hybridization of four tyrosinase candidates. Abbreviations: DT - digestive tract; he – cement gland with heterogeneous granules; ho – cement gland with homogeneous granules.

## Discussion

### Diversity of tyrosinases potentially involved in adhesive protein maturation

Various marine invertebrates rely on quinone-tanned biomaterials as durable glues or protective varnishes (Waite, 1990). Two well-studied examples of such materials are the byssus produced by mussels and the cement secreted by tube-building worms (Waite, 1990). These materials contain a catechol known as DOPA, or 3,4-dihydroxyphenylalanine, a modified amino acid which is widely present in nature. DOPA is essential for bonding with the substrate and providing cohesive cross-links within the adhesive materials of these organisms. The enzymes responsible for its production are the tyrosinases which play a crucial role in various biological functions. Although DOPA has been extensively studied in marine adhesion, the specific enzymes responsible for its production are still not well understood. However, several studies have highlighted the fact that not one but several different tyrosinases may be involved in the maturation of adhesive proteins in a single organism (Guerette *et al*. 2013; Qin *et al*. 2016; Buffet *et al*. 2018). In this study, our first objective was to identify the tyrosinase sequences involved in the adhesive systems of mussels and tubeworms by comparing a set of reference sequences known to be present in the adhesive secretions of some species with the transcriptomes of our two model species, the blue mussel *M. edulis* and the honeycomb worm *S. alveolata*. For both species, more than 80 different tyrosinase-like sequences were retrieved by the BLAST searches, but these numbers were greatly reduced when only (almost) full-length proteins with a tyrosinase domain were considered. Moreover, differential expression, mass spectrometry analyses and *in situ* hybridisation experiments were used to pinpoint the candidate expressed in adhesive glands and secreted.

In the blue mussel, 17 candidates were identified by the *in silico* analyses, among which 7 were shown to be present in the byssus and/or in the foot tissues by mass spectrometry analyses. In particular, 5 tyrosinases were detected in induced byssal threads with a high peptide coverage (Medu TYR1 and 2, and Medu-TYR10 to 12). The mRNAs coding for these 5 proteins, but also those coding for 3 other candidates sharing high homology with reference tyrosinases from *M. coruscus* and *P. viridis*, were localised in the foot glands. Two patterns of expression were observed: three candidates (Medu-TYR10 to 12) are each specific in a single gland type while the other five (Medu-TYR1, 2, 4, 5 and 15) have been localised in different glands (Fig. 6). The gland-specific mRNA expression of these different tyrosinases corresponds to the localisation of the corresponding protein in the foot (distal half, proximal half, or both) for candidates which have been detected in mass spectrometry (Table 1; Fig. 6). It also matches the distribution of their homologues within the byssus of *M. coruscus* (Table 1). For instance, Medu-TYR12, whose mRNA is specifically expressed in the plaque gland, is the closest homologue to a tyrosinase-like protein (encoded by cDNA KP876481; Table 1) identified in the byssal plaque in *M. coruscus* (Qin *et al*., 2016). It is not surprising to find tyrosinases in the three different foot glands involved in byssal threads synthesis as they all produce DOPA-containing proteins (Waite, 2017). The plaque gland is known to contain more than ten different proteins (DeMartini *et al*., 2017), including mfp-3 and -5 which are the byssal proteins with the highest DOPA content (20 and 30 mol%, respectively) (Papov *et al*., 1995; Waite & Qin, 2001). The cuticle gland produces mfp-1, which contains 15 mol% of DOPA, as well as handful of other, less-characterized proteins (Waite & Tanzer, 1981; DeMartini *et al*., 2017). Finally, although the preCols and thread matrix proteins forming the core of byssal thread contain lower amounts of DOPA (Qin *et al*., 1997; Sagert & Waite, 2009), it plays a critical role in thread core cross-linking (Priemel *et al*., 2020), and thus, this modified amino acid is also present in the core gland.

**Figure 6.**
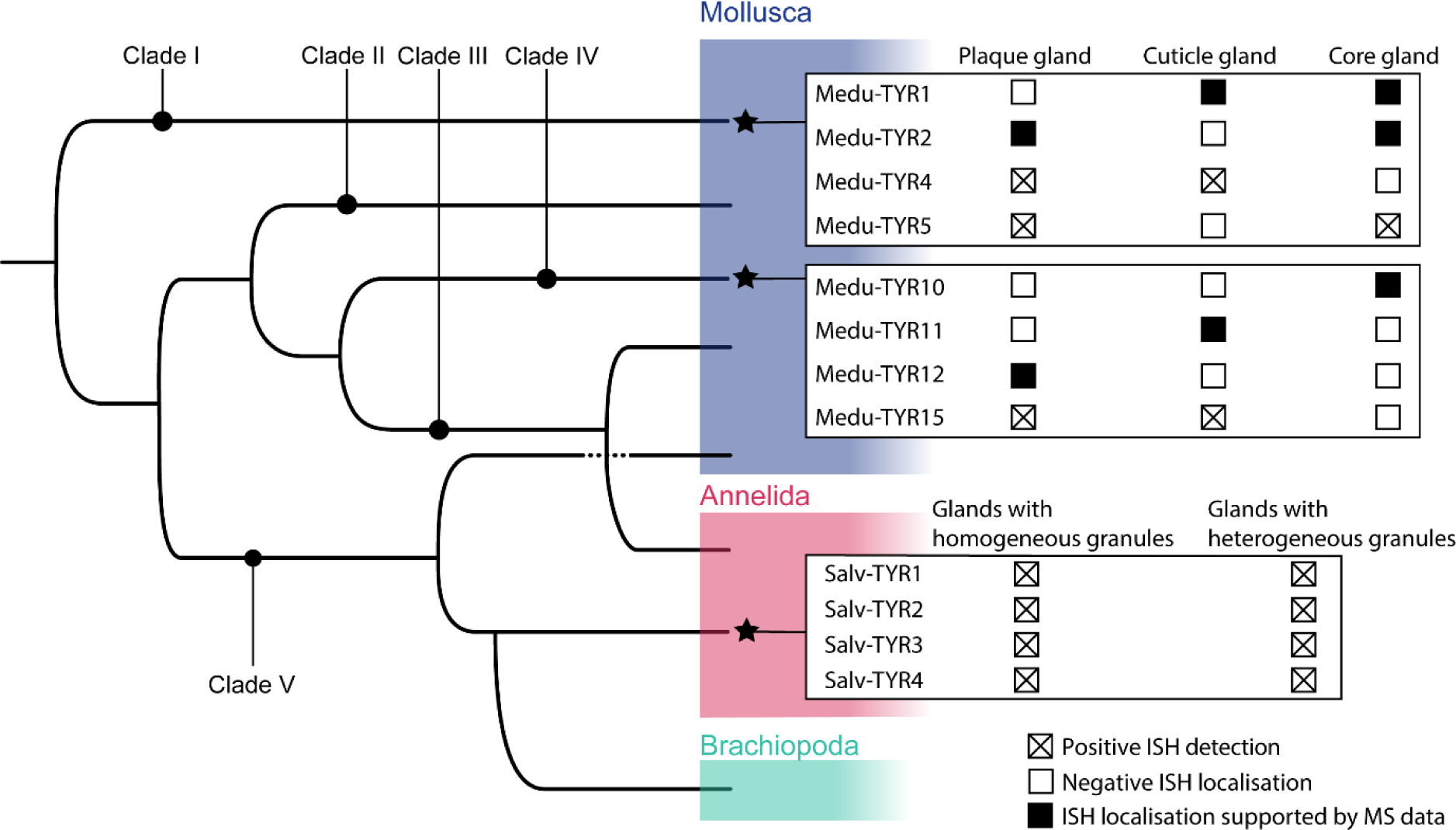
Tyrosinases involved in the adhesive systems of the mussel *Mytilus edulis* and the tubeworm *Sabellaria alveolata*. Simplified tyrosinase phylogenetic tree in which stars indicate the clades comprising enzymes expressed in adhesive glands. The gland specificity of each tyrosinase, as highlighted by *in situ* hybridisation experiments, is illustrated on the right. In the mussel, the ISH localisation of several candidates is supported by mass spectrometry data even though the experimental protocol used (foot distal part vs foot proximal part) does not allow a precise gland assignment.

In the honeycomb worm, 28 candidate tyrosinase transcripts were retrieved from the transcriptomic analysis. The comparison of these sequences with the differential transcriptome of Buffet *et al*. (2018) for the same species allowed us to reduce the list to 13 candidates. These transcripts are highly overexpressed in the parathorax, the region of the worm containing the cement glands. A proteomic analysis was conducted on tubes reconstructed by the worm using glass beads but, unfortunately, no tyrosinase or cement protein was detected, suggesting that the proteins could not be extracted from the cement spots binding the beads. Four transcripts, among the most differentially expressed and showing the highest homology with the cDNA encoding the reference tyrosinase from the sandcastle worm *P. californica* (JN607213.2), were selected for localisation by *in situ* hybridization. These tyrosinase mRNAs were expressed in both types of unicellular cement glands, the cells with homogeneous granules and the cells with heterogeneous granules, similarly to what was observed in *P. californica* (Wang & Stewart, 2012). There was therefore no gland specificity for the four candidates we selected. In *P. californica*, at least two adhesive proteins, Pc-1 and Pc-2, are known to contain DOPA (about 10 and 7%, respectively; Waite *et al*., 1992). The former is produced by cement glands with heterogeneous granules and the latter by glands with homogeneous granules (Wang & Stewart, 2012). Their homologues, Sa-1 and Sa-2, have been identified in *S. alveolata* (Becker *et al*., 2012) and preliminary results suggest that their distribution in the cement glands corresponds to the one in *P. californica* (unpubl. obs.). Thus, in the honeycomb worm too, each type of cement gland would produce at least one DOPA-rich protein, which would explain the expression of tyrosinases in both glands.

Although the high diversity of tyrosinase candidates in both model species is suggestive of a variety of functions, linking their sequences with their different functionalities (monooxygenase vs oxidase) may prove difficult. Tyrosinases and catechol oxidases share a very similar active site architecture in which three distinct oxidation states have been identified, oxy-, deoxy-, and met-states, based on the structure of the bicopper structure of the active centre (Ramsden & Riley, 2014). Transitions from one state to another lead to the molecular mechanisms involved in the monophenolase or diphenolase catalytic activity (Ramsden & Riley, 2014). However, some enzymes appear to show only the oxidase activity (Pretzler & Rompel, 2018). Moreover, while the active site of tyrosinases is highly conserved, variations exist in their sequences, size, glycosylation, and activation (Jaenicke & Decker, 2003). To date, the structural motifs responsible for the different activities remain elusive (Pretzler & Rompel, 2018). Only homology with enzymes of known function or expression pattern could therefore be used to investigate the function of the new tyrosinase candidates from *M. edulis* and *S. alveolata*.

In mussels, DOPA plays important interfacial adhesive and bulk cohesive roles during byssus fabrication (Lee *et al*. 2011, Waite 2017). Priemel *et al*. (2020) have identified two distinct DOPA-based cross-linking pathways, one achieved by oxidative covalent cross-linking and the other by the formation of metal coordination interactions under reducing conditions. The former takes place in the thread core while the latter occurs in the cuticle, plaque foam and plaque-surface interface (Priemel *et al*., 2020). In the core gland, preCol-D was shown to possess several DOPA residues at the N-terminus (Qin *et al*., 1997) which, after secretion, are oxidised into dopaquinone for covalent cross-link formation (Priemel *et al*., 2020). It is therefore tempting to postulate that the core gland-specific tyrosinase, Medu-TYR12, which is also one of the most highly expressed, could have a catechol oxidase activity. A catechol oxidase was extracted from the foot and byssus of *M. edulis* by Waite (1985), but it is not possible to link it to any of our candidate as no sequence was reported. On the other hand, the DOPA-rich proteins from the cuticle and plaque glands should be prevented from oxidation after secretion to promote DOPA-metal coordination intermolecular bonds (Schmitt *et al*., 2015; Priemel *et al*., 2020). This is possible thanks to a locally low pH and the presence of reducing conditions, often from the sulfhydryl groups of cysteine-rich proteins such as mfp-6 (Yu *et al*., 2011). Following a similar reasoning as above, Medu-TYR11 and 12, which are specific to the cuticle and plaque gland, respectively, could then only display a monophenolase activity. Unexpectedly, Medu-TYR15, whose corresponding transcript is expressed in both the plaque and cuticle glands, is the closest homologue to a catechol oxidase from *M. galloprovincialis* (Table 1). However, interestingly, this catechol oxidase, when expressed recombinantly in bacteria, showed no catalytic activity but, instead, exhibited a thiol-dependent antioxidant activity suppressing DOPA oxidation (Wang *et al*., 2019). Medu-TYR15 was not detected in the byssus or the foot by our proteomic analyses, however.

In tubeworms, it was demonstrated that the tyrosinase identified in *P. californica* catalyses only catechol oxidation. This observation was made through substrates and inhibitors profiling on the isolated secretory granules and secreted cement (Wang & Stewart, 2013). As Salv-TYR1 to 4 are close homologues to this catechol oxidase and display the same expression pattern in both cement glands, it is likely that they would also show a diphenolase activity. In *P. californica*, the oxidation of DOPA to dopaquinone leads to the formation of cysteinyl-DOPA covalent cross-links within the cement (Zhao *et al*., 2005).

### Evolution of tyrosinases in Lophotrochozoa

The high number and diversity of tyrosinases identified in *M. edulis* and *P. californica* prompted us to examine the phylogenetic relationship they have together and with other lophotrochozoan tyrosinases.

Firstly, we built a cluster map of all spiralian sequences containing a tyrosinase domain retrieved from the NCBI nr database. Tyrosinase sequences from Platyhelminthes formed a cluster clearly separated from another large cluster comprising sequences from Brachiopoda, Annelida, and Mollusca. This reflects the evolution of type-3 copper proteins in animals in which three ancestral subclasses (α, β and γ) have been described, with differential losses or expansions of one or more of these subclasses in specific phyla (Aguilera *et al*., 2013). Platyhelminthes possess γ-subclass, transmembrane tyrosinases whereas the three other phyla possess α-subclass, secreted tyrosinases (Aguilera *et al*., 2013). Molluscan hemocyanins also belong to the α-subclass (Aguilera *et al*., 2013), but form a well-separated cluster compared to tyrosinases.

Secondly, a phylogenetic tree was constructed to examine the relationships between lophotrochozoan tyrosinases, using molluscan hemocyanins as the outgroup. The tyrosinase tree is fairly complex, with five main clades that do not necessarily reflect the main taxonomic groups. Mollusc, annelid and brachiopod tyrosinases appear to be paraphyletic and their sequence variation would not be solely influenced by the evolutionary distance between lineages. Indeed, three of the five clades in the tree we generated (clades I, II and IV) contain exclusively sequences from bivalves while the other two clades (III and V) gather sequences from different phyla or mollusc classes. In their study about the evolution of the tyrosinase gene family in bivalve molluscs, Aguilera *et al*. (2014) reported that tyrosinase sequences were separated into two clades: an ancestral clade (A) comprising tyrosinases from all mollusc classes and a bivalve-specific clade (B). Based on the sequences shared by the two studies, our clades I, II, III and V would correspond to their clade A, and our clade IV to their clade B. The different tree topology we obtained could be linked to the inclusion of annelid and brachiopod sequences in our tree, a different trimming of the sequences, or a different rooting of the tree. It should also be noted some of the basal nodes in our tree are poorly supported.

The high number of transcripts obtained during our *in silico* analyses supports the hypothesis that the tyrosinase gene family has undergone large independent expansions in bivalve molluscs (Aguilera *et al*., 2014) as well as in sabellariid polychaetes (Buffet *et al*., 2018). This apparently also includes tyrosinases potentially involved in the maturation of adhesive proteins, with 6 and 4 enzymes identified in *M. edulis* and *S. alveolata*, respectively, as shown by proteomic data and/or *in situ* hybridization results (Fig. 6). Mussel byssus and tubeworm cement tyrosinase sequences are separated in distinct clusters. This suggests an independent functional evolution of tyrosinases involved in the maturation of bioadhesives in these two lineages. Interestingly, the mussel tyrosinases are also distributed in two different clusters (I and IV) and enzymes from both clusters are co-expressed in a same foot gland. The identification of byssal tyrosinases in these two different clusters is congruent with the results of Aguilera *et al*. (2014) indicating an early gene duplication in bivalve evolution. These genes would then have undergone further independent duplication and divergence to acquire new gene functions.

## Conclusion

Honeycomb worms and blue mussels are frequently found together in the intertidal zone, where they encounter similar environmental challenges. Both organisms rely on a DOPA-based adhesive system and our study has identified a catalogue of tyrosinase enzymes involved in the maturation of adhesive proteins in *M. edulis* and *S. alveolata*. The diversity of tyrosinases highlighted in the two species suggests the coexistence of different functions (monophenol monooxygenase or catechol oxidase activity) or different substrate specificities. However, the exact role of the different enzymes needs to be further investigated. Phylogenetic analyses support the hypothesis of independent expansion and parallel evolution of tyrosinases involved in adhesive protein maturation in both lineages, supporting the convergent evolution of their DOPA-based adhesion. These results contribute to our understanding of the molecular basis of adhesion mechanisms in marine organisms, but also pave the way towards the use of specific tyrosinases in development of biomimetic adhesives.

## Methods

### Animal collection and maintenance

Individuals of *Mytilus edulis* Linnaeus, 1758, were collected intertidally in Audresselles (Pas-de-Calais, France). Reef fragments of *Sabellaria alveolata* Linnaeus, 1767, were collected at low tide on the “plage de Ris’’ at Douarnenez, France (48°05’34.6’’N 4°17’55.0’’W), or were obtained from the Station Biologique de Roscoff (Finistere, France). All animals were then transported to the Laboratory of Biology of Marine Organisms and Biomimetics (University of Mons, Belgium), where they were kept in a re-circulating aquarium (13°C, 33 psu salinity).

### Transcriptome sequencing and *de novo* assembly

In the present study, two tissue transcriptomes have been generated: one from the foot of the blue mussel *M. edulis* and one from the anterior part of the honeycomb worm *S. alveolata*.

RNA extraction, library construction and sequencing were performed at the GIGA Genomics platform (Liège, Belgium). After dissection, mussel feet and tubeworm anterior parts (corresponding to the head and parathoracic region) were immediately frozen with liquid nitrogen and stored at −80°C until use. Total RNA was extracted from 100 mg of frozen tissue using Trizol (Life Technologies, Carlsbad, CA) and its quality was assessed using the Bioanalyser 2100 (Agilent). Truseq Stranded mRNA Sample Preparation kit (San Diego, CA) was used to prepare a library from 500 ng of total RNA. Poly-adenylated RNAs were purified with oligo (dT)-coated magnetic beads (Sera-Mag Magnetic Oligo(dT) beads, Illumina) and then chemically fragmented to a length of 100 to 400 nucleotides -with a majority of the fragments at about 200 bp (base pairs)-by using divalent cations at 94°C for 5 minutes. These short fragments were used as a template for reverse-transcription using random hexamers to synthetize cDNA, followed by end reparation and adaptor ligation according to the manufacturer’s protocol (Illumina, San Diego, CA). Finally, the ligated library fragments were purified and enriched by solid-phase PCR following Illumina’s protocol. The library quality was validated on the Bioanalyser 2100. The high-throughput sequencing was conducted by a HiSeq 2000 platform (Illumina, San Diego, CA) to obtain 2×100-bp paired-end reads according to manufacturer’s instructions. Real-time quality control was performed to ensure that most read quality score was higher than 30. The raw sequencing data have been deposited in the NCBI Sequence Read Archive with accession numbers PRJNA1125179 and PRJNA1125180 for *M. edulis* and *S. alveolata*, respectively.

Transcriptome quality was checked using Fast QC software (Babraham Bioinformatics). The Trinity software suite (Grabherr *et al*., 2011) which comprises a quality filtering function was used with default parameters to *de novo* assemble the raw reads with overlapping nucleic acid sequence into contigs. Transcriptome completeness was evaluated using BUSCO (v3.0.2) analyses on assembled transcripts (Waterhouse *et al*., 2018). Scores were calculated using Metazoan_odb10 lineage data.

### *In silico* tyrosinase sequences identification

To find the tyrosinase sequences putatively involved in adhesive protein maturation, tBLASTn searches were performed on the two transcriptomes using a reference dataset (see Suppl. Table 1) consisting of byssus-specific tyrosinase sequences found in *M. coruscus* (Qin *et al*., 2016) and *P. viridis* (Guerette *et al*., 2013) and of a cement tyrosinase sequence from the tubeworm *P. californica* (Wang & Stewart, 2012). Several transcripts encoding tyrosinase-like proteins were obtained for both species. They were subsequently used in a reciprocal tBLASTn search against the NCBI non-redundant protein database and only those having a tyrosinase as the best-hit match were kept for further analyses.

The selected transcripts were translated *in silico* (Expasy, Translate tool) and analyzed by looking for the tyrosinase copper-binding domain (IPR002227) using InterPro (https://www.ebi.ac.uk/interpro/search/sequence/) (Paysan-Lafosse *et al*., 2023) and for the presence of a signal peptide using SignalP-6.0 (https://services.healthtech.dtu.dk/service.php?SignalP) (Teufel *et al*., 2022).

### Protein extraction and mass spectrometry analyses

To assess whether tyrosinase enzymes are secreted in the adhesive materials of mussels and tubeworms, proteomic analyses were conducted on induced byssal threads and reconstructed tubes.

For *M. edulis*, the secretion of fresh byssal threads was induced by injecting a 0.56 M solution of KCl at the base of the foot as described by Tamarin *et al*. (Tamarin *et al*., 1976). For protein extraction, approximately 30 byssal threads were crushed in 500 µL of a solution containing 5% acetic acid and 8 M urea (Rzepecki *et al*., 1992). The extract was then centrifuged for 20 min at 16,000g and the supernatant containing the proteins was collected. For *S. alveolata*, single tubes with their dwelling worm were isolated from the reef fragment. The upper third of each tube was then removed and the worms were allowed to reconstruct this missing portion of their tube with glass beads (425-600 µm in diameter; Sigma) (Jensen & Morse, 1988). Approximately 500 mg of freshly rebuilt tube fragments were subjected to protein extraction using a 1.5 M Tris–HCl buffer (pH 8.5) containing 7 M guanidine hydrochloride (GuHCl), 20 mM ethylenediaminetetraacetate (EDTA), and 0.5 M dithiothreitol (DTT) (Tris-GuHCl buffer). This mixture was incubated for 1 hour at 60°C under agitation and then centrifuged as described above.

To further process proteins, both samples were treated with a 2.5-fold excess (w/w) of iodoacetamide to dithiothreitol (DTT), for 20 minutes in the dark at room temperature, to carbamidomethylate the sulfhydryl groups. The reaction was then stopped by adding an equal quantity of β-mercaptoethanol to iodoacetamide. The extract was centrifuged at 13,000 rpm for 15 minutes at 4°C and the supernatant was collected. The protein concentration was determined using the Non-Interfering Protein Assay Kit (Calbiochem, Darmstadt, Germany) with bovine serum albumin as a protein standard. For each sample, 50 μg of proteins were then precipitated overnight at −20°C in 80% acetone. After a 15-minute centrifugation at 13,000 rpm and evaporation of acetone, the resulting pellet was subjected to overnight enzymatic digestion using modified porcine trypsin at an enzyme/substrate ratio of 1/50, at 37°C in 25 mM NH_4_HCO_3_. The reaction was stopped by adding formic acid to a final concentration of 0.1% (v/v) (Hennebert *et al*., 2015). Tryptic peptides were analysed by LC connected to a hybrid quadrupole time-of-flight TripleTOF 6600 mass spectrometer (AB SCIEX, Concord, ON). Byssal threads and reconstructed tubes MS/MS data were searched for protein candidates against a database composed of the six open reading frames (ORFs) of the transcriptome of the foot of *M. edulis* or of the transcriptome of the anterior part of *S. alveolata*, respectively, using the Protein Pilot software (version 5.0.1). The samples with a false discovery rate (FDR) above 1.0% were excluded from subsequent analyses.

A proteomic analysis was also carried out on the mussel foot tissues. Feet were dissected and cut transversely into two halves to separate the proximal part from the distal part. The samples were homogenized in a Potter-Elvehjem tissue grinder with a 4% solution of SDS in Tris-HCl buffer (pH 7.4), with protease inhibitors (EDTA-free), and 5 units of DNase per 100 mg of foot tissue. The samples were then sonicated three times for 30 seconds each and left at room temperature for 30 minutes, followed by overnight incubation at 4°C. The protein content was quantified using an RCDC kit (Biorad). Aliquots of 15 µg of proteins were reduced and alkylated. The samples were then treated with the 2-D Clean-Up Kit (GE Healthcare) to eliminate impurities not compatible with mass spectrometry analyses and recovered in 50 mM ammonium bicarbonate buffer. Trypsin digestion was carried out for 16 hours at an enzyme/substrate ratio of 1/50 at 37°C, the reaction was stopped by adding trifluoroacetic acid, and the samples were dried using a speed vac. The tryptic peptides were dissolved in water with 0.1% trifluoroacetic acid and purified using a Zip-Tip C18 High Capacity. They were analysed by reverse-phase HPLC–ESI-MS/MS using a nano-UPLC (nanoAcquity, Waters) connected to an ESI-Q-Orbitrap mass spectrometer (Q Extractive Thermo) in positive ion mode. MS/MS data were analysed against a database composed of the six open reading frames (ORFs) of the mussel foot transcriptome using the MaxQuant software (version 1.4.1.2). The peptide mass tolerance was set to ±10 ppm, and fragment mass tolerance was set to ±0.1 Da. The oxidation of the amino acids tyrosine, arginine and proline, as well as the carbamidomethylation of cysteine and the phosphorylation of serine were defined as fixed modifications, and the oxidation of methionine as variable modifications. Protein identifications were considered significant if proteins are identified with at least two peptides per protein taking into account only an FDR<0.01.

### Localization of tyrosinase mRNA

The best tyrosinase sequence candidates found in the *in silico* and proteomic analyses were localized using in *situ* hybridization (ISH) technique to see if they are well expressed in the adhesive glands of both studied species. RNA was extracted from three parathoracic parts of honeycomb worms and three blue mussel feet using TRIzol^TM^ Reagent kit (Thermofisher). The cDNA synthesis from the RNA extracted was done using Reverse transcription kit, Roche. Later, double-stranded DNA templates were amplified by PCR using PCR Using Q5 High-Fidelity DNA Polymerase kit method, with primer designed by Open Primer 3 (bioinfo.ut.ee/primer3/) with an optimal probe length between 700 and 900 bp. A second PCR was done with T7 promoter binding site (5’-GGATCCTAATACGACTCACTATAGG-3’) added to reverse strand PCR primers. PCR products were purified using the Wizard SV Gel and PCR clean-up system kit (Promega) and used for RNA probe synthesis. Digoxigenin (DIG)-labelled RNA probes were then synthesized with the kit DIG RNA Labelling Kit (Roche) with T7 RNA polymerase and DIG–dUTP. In situ hybridization was performed according to Lengerer *et al*., 2018. The probes were used on parathoracic sections showing the adhesive glands for the honeycomb worm, and transversal sections of the blue mussel foot, at a concentration of 0.2ng/µl and detected with antidigoxigenin-AP Fab fragments (Roche) at a dilution of 1:2000. The signal was developed using the NBT/BCIP system (Roche) at a dilution of 1:50 at 37°C. Sections were observed using a Zeiss Axioscope A1 microscope connected to a Zeiss AxioCam 305 color camera.

To confirm the localization of the candidates in the blue mussel, we conducted a complementary whole mount *in situ* hybridization. For each candidate, the mussel feet were divided into two parts along the frontal axis. The protocol was done in 6-well plates according to Pfister *et al*., 2007, except for these steps; the *in situ* hybridization was carried out on samples re-incubated at 55°C for 72 hours, and color development was performed in the dark at 37°C using an NBT/BCIP system (Roth) until a satisfactory precipitate coloration was achieved.

### Phylogenetic analyses and clustering analyses of tyrosinase sequences

To gain a deeper understanding of the relationship between all the tyrosinase enzymes, we conducted a CLANS analysis on a comprehensive dataset. This dataset included lophotrochozoan sequences in FASTA format, each containing a pfam00264 domain. These sequences were retrieved from the NCBI database (retrieved on September 20, 2022). In addition, we also incorporated the previously identified sequences obtained through *in silico* and proteomic analyses. We also incorporated three tyrosinase transcripts from the transcriptome of *Mytilus edulis*, which exhibited higher expression levels in mantle tissues (Matthew J. Harrington personal communication). To expand the diversity of tubeworm tyrosinases within the phylum Annelida, we obtained transcriptomes from *Phragmatopoma caudata* (Buffet *et al*., 2018). We conducted a local tBLASTn search targeting tyrosinase mRNA sequences that may play a role in protein cement maturation, using the previously mentioned *P. californica* sequence. As the number of sequences was relatively limited for Annelida compared with other phyla, we conducted a complementary analysis. In this analysis, we searched for the pfam00264 domain among the sequences found in the transcriptomes of *Capitella teleta* (3 transcripts), *Owenia fusiformis* (13 transcripts), *Oasisia alvinae* (4 transcripts), and *Riftia pachyptila* (3 transcripts), all of which are available on https://github.com/ChemaMD. The CLANS analysis was based on all-against-all sequence similarity using BLAST searches with the BLOSUM62 matrix (https://toolkit.tuebingen.mpg.de/clans/) (Frickey & Lupas, 2004).

We performed a phylogenetic analysis using the tyrosinase sequences from two closely clustered groups identified in the CLANS analysis, and which contained the sequences from the two studied species. First, we conducted a multiple alignment with all these sequences, totaling 312 sequences, using the MAFFT algorithm (using the automated parameters of Mafft v7.490 implemented on Geneious Prime 2023.2.1, Katoh *et al*., 2019). Subsequently, we trimmed this alignment using the online TrimAL tool implemented on Phylemon2 (Capella-Gutierrez *et al*., 2009) with the Automated1 option parameter. We selected two tyrosinase sequences from Mollusca, which contained a hemocyanin cluster as outgroup. The construction of the phylogenetic tree was carried out using the IQ-TREE (1.6.12) software, using the maximum likelihood method and performing ultrafast bootstrap (UFBoot) analysis with 1000 replicates (Hoang *et al*., 2018). According to the software’s BIC scores, the best-fit model was determined to be the WAG+F+I+G4 model (Trifinopoulos *et al*., 2016). The tree was modified using the software iTOL (version 6.9) (Letunic & Bork, 2021).

## Acknowledgements

We thank Dominique Baiwir from the GIGA Proteomics Facility (ULiège, Belgium) for the mass spectrometry-based analyses conducted on mussel feet, as well as Antoine Flandroit, Nathan Puozzo and Paolo Rosa for their help with microphotography and figures. This work was supported by the Fund for Scientific Research of Belgium (F.R.S.-FNRS) through a FRIA doctoral grant to E.D., a CR postdoctoral fellowship to B.M., and a “Projet de Recherche” (T.0088.20). P.F. is Research Director of the F.R.S.-FNRS.

## References

Aguilera, F., McDougall, C., & Degnan, B. M. (2013). Origin, evolution and classification of type-3 copper proteins: Lineage-specific gene expansions and losses across the Metazoa. BMC Evolutionary Biology, 13(1), 96.

Aguilera, F., McDougall, C., & Degnan, B. M. (2014). Evolution of the tyrosinase gene family in bivalve molluscs: Independent expansion of the mantle gene repertoire. Acta Biomaterialia, 10(9), 3855–3865.

Almeida, M., Reis, R. L., & Silva, T. H. (2020). Marine invertebrates are a source of bioadhesives with biomimetic interest. Materials Science and Engineering: C, 108, 110467.

Andersen, S. O. (2010). Insect cuticular sclerotization: A review. Insect Biochemistry and Molecular Biology, 40(3), 166–178.

Becker, P. T., Lambert, A., Lejeune, A., Lanterbecq, D., & Flammang, P. (2012). Identification, Characterization, and Expression Levels of Putative Adhesive Proteins From the Tube-Dwelling Polychaete *Sabellaria alveolata*. The Biological Bulletin, 223(2), 217–225.

Buffet, J.-P., Corre, E., Duvernois-Berthet, E., Fournier, J., & Lopez, P. J. (2018). Adhesive gland transcriptomics uncovers a diversity of genes involved in glue formation in marine tube-building polychaetes. Acta Biomaterialia, 72, 316–328.

Burmester, T., & Schellen, K. (1996). Common origin of arthropod tyrosinase, arthropod hemocyanin, insect hexamerin, and dipteran arylphorin receptor. Journal of Molecular Evolution, 42(6), 713–728.

Capella-Gutierrez, S., Silla-Martinez, J. M., & Gabaldon, T. (2009). trimAl: A tool for automated alignment trimming in large-scale phylogenetic analyses. Bioinformatics, 25(15), 1972–1973.

Davey, P. A., Power, A. M., Santos, R., Bertemes, P., Ladurner, P., Palmowski, P., Clarke, J., Flammang, P., Lengerer, B., Hennebert, E., Rothbächer, U., Pjeta, R., Wunderer, J., Zurovec, M., & Aldred, N. (2021). Omics-based molecular analyses of adhesion by aquatic invertebrates. Biological Reviews, 96(3), 1051–1075.

Del Marmol, V., & Beermann, F. (1996). Tyrosinase and related proteins in mammalian pigmentation. FEBS Letters, 381(3), 165–168.

Delroisse, J., Kang, V., Gouveneaux, A., Santos, R., & Flammang, P. (2023). Convergent Evolution of Attachment Mechanisms in Aquatic Animals. In V. L. Bels & A. P. Russell (Éds.), Convergent Evolution (p. 523–557). Springer International Publishing.

DeMartini, D. G., Errico, J. M., Sjoestroem, S., Fenster, A., & Waite, J. H. (2017). A cohort of new adhesive proteins identified from transcriptomic analysis of mussel foot glands. Journal of The Royal Society Interface, 14(131), 20170151.

Dennell, R. (1958). The hardening of insect cuticles. Biological Reviews, 33(2), 178–196.

Drexel, R., Siegmund, S., Schneider, H.-J., Linzen, B., Gielens, C., Préaux, G., Lontie, R., Kellermann, J., & Lottspeich, F. (1987). Complete Amino-Acid Sequence of a Functional Unit from a Molluscan Hemocyanin (*Helix pomatia*). Biological Chemistry Hoppe-Seyler, 368(1), 617–636.

Endrizzi, B. J., & Stewart, R. J. (2009). Glueomics : An Expression Survey of the Adhesive Gland of the Sandcastle Worm. The Journal of Adhesion, 85(8), 546–559.

Esposito, R., D’Aniello, S., Squarzoni, P., Pezzotti, M. R., Ristoratore, F., & Spagnuolo, A. (2012). New Insights into the Evolution of Metazoan Tyrosinase Gene Family. PLoS ONE, 7(4), e35731.

Flammang, P., Lambert, A., Bailly, P., & Hennebert, E. (2009). Polyphosphoprotein-Containing Marine Adhesives. The Journal of Adhesion, 85(8), 447–464.

Frickey, T., & Lupas, A. (2004). CLANS: A Java application for visualizing protein families based on pairwise similarity. Bioinformatics, 20(18), 3702–3704.

González-Santoyo, I., & Córdoba-Aguilar, A. (2012). Phenoloxidase : A key component of the insect immune system: Biochemical and evolutionary ecology of PO. Entomologia Experimentalis et Applicata, 142(1), 1–16.

Grabherr, M. G., Haas, B. J., Yassour, M., Levin, J. Z., Thompson, D. A., Amit, I., Adiconis, X., Fan, L., Raychowdhury, R., Zeng, Q., Chen, Z., Mauceli, E., Hacohen, N., Gnirke, A., Rhind, N., Di Palma, F., Birren, B. W., Nusbaum, C., Lindblad-Toh, K., … Regev, A. (2011). Full-length transcriptome assembly from RNA-Seq data without a reference genome. Nature Biotechnology, 29(7), 644–652.

Guerette, P. A., Hoon, S., Seow, Y., Raida, M., Masic, A., Wong, F. T., Ho, V. H. B., Kong, K. W., Demirel, M. C., Pena-Francesch, A., Amini, S., Tay, G. Z., Ding, D., & Miserez, A. (2013). Accelerating the design of biomimetic materials by integrating RNA-seq with proteomics and materials science. Nature Biotechnology, 31(10), 908–915.

Hellio, C., Bourgougnon, N., & Gal, Y. L. (2000). Phenoloxidase (E.C. 1.14.18.1) from the byssus gland of *Mytilus edulis*: Purification, partial characterization and application for screening products with potential antifouling activities. Biofouling, 16(2-4), 235–244.

Hennebert, E., Leroy, B., Wattiez, R., & Ladurner, P. (2015). An integrated transcriptomic and proteomic analysis of sea star epidermal secretions identifies proteins involved in defense and adhesion. Journal of Proteomics, 128, 83–91.

Hoang, D. T., Chernomor, O., Von Haeseler, A., Minh, B. Q., & Vinh, L. S. (2018). UFBoot2 : Improving the Ultrafast Bootstrap Approximation. Molecular Biology and Evolution, 35(2), 518–522.

Hofman, A. H., van Hees, I. A., Yang, J., & Kamperman, M. (2018). Bioinspired Underwater Adhesives by Using the Supramolecular Toolbox. Advanced Materials, 30(19), 1704640.

Huan, P., Liu, G., Wang, H., & Liu, B. (2013). Identification of a tyrosinase gene potentially involved in early larval shell biogenesis of the Pacific oyster *Crassostrea gigas*. Development Genes and Evolution, 223(6), 389–394.

Jaenicke, E., & Decker, H. (2003). Tyrosinases from crustaceans form hexamers. Biochemical Journal, 371(2), 515–523.

Jensen, R. and Morse, D. (1988). The bioadhesive of *Phragmatopoma californica* tubes: a silk-like cement containing L-DOPA. J Comp Physiol B. 158:317–324.

Kamino, K. (2010). Molecular Design of Barnacle Cement in Comparison with Those of Mussel and Tubeworm. The Journal of Adhesion, 86(1), 96–110.

Katoh, K., Rozewicki, J., & Yamada, K. D. (2019). MAFFT online service : Multiple sequence alignment, interactive sequence choice and visualization. Briefings in Bioinformatics, 20(4), 1160–1166.

Laumer, C. E., Bekkouche, N., Kerbl, A., Goetz, F., Neves, R. C., Sørensen, M. V., Kristensen, R. M., Hejnol, A., Dunn, C. W., Giribet, G., & Worsaae, K. (2015). Spiralian Phylogeny Informs the Evolution of Microscopic Lineages. Current Biology, 25(15), 2000–2006.

Lee, H., Scherer, N. F., & Messersmith, P. B. (2006). Single-molecule mechanics of mussel adhesion. Proceedings of the National Academy of Sciences, 103(35), 12999–13003.

Lengerer, B., Wunderer, J., Pjeta, R., Carta, G., Kao, D., Aboobaker, A., Beisel, C., Berezikov, E., Salvenmoser, W., & Ladurner, P. (2018). Organ specific gene expression in the regenerating tail of Macrostomum lignano. Developmental Biology, 433(2), 448–460.

Letunic, I., & Bork, P. (2021). Interactive Tree Of Life (iTOL) v5 : An online tool for phylogenetic tree display and annotation. Nucleic Acids Research, 49(W1), W293–W296.

Li, X., Li, S., Huang, X., Chen, Y., Cheng, J., & Zhan, A. (2021). Protein-mediated bioadhesion in marine organisms : A review. Marine Environmental Research, 170, 105409.

Papov, V. V., Diamond, T. V., Biemann, K., & Waite, J. H. (1995). Hydroxyarginine-containing Polyphenolic Proteins in the Adhesive Plaques of the Marine Mussel *Mytilus edulis*. Journal of Biological Chemistry, 270(34), 20183–20192.

Paysan-Lafosse, T., Blum, M., Chuguransky, S., Grego, T., Pinto, B. L., Salazar, G. A., Bileschi, M. L., Bork, P., Bridge, A., Colwell, L., Gough, J., Haft, D. H., Letunić, I., Marchler-Bauer, A., Mi, H., Natale, D. A., Orengo, C. A., Pandurangan, A. P., Rivoire, C., … Bateman, A. (2023). InterPro in 2022. Nucleic Acids Research, 51(D1), D418–D427.

Pearson, W.R. (2013). An introduction to sequence similarity (“homology”) searching. Current Protocols in Bioinformatics, Chapter 3, Unit3.1.

Petrone, L. (2013). Molecular surface chemistry in marine bioadhesion. Advances in Colloid and Interface Science, 195-196, 1–18.

Pfister, D., De Mulder, K., Philipp, I., Kuales, G., Hrouda, M., Eichberger, P., Borgonie, G., Hartenstein, V., & Ladurner, P. (2007). The exceptional stem cell system of *Macrostomum lignano* : Screening for gene expression and studying cell proliferation by hydroxyurea treatment and irradiation. Frontiers in Zoology, 4(1), 9.

Polivares, C., & Solano, F. (2009). New insights into the active site structure and catalytic mechanism of tyrosinase and its related proteins. Pigment Cell & Melanoma Research, 22(6), 750–760.

Pretzler, M., & Rompel, A. (2018). What causes the different functionality in type-III-copper enzymes? A state of the art perspective. Inorganica Chimica Acta, 481, 25–31.

Priemel, T., Degtyar, E., Dean, M. N., & Harrington, M. J. (2017). Rapid self-assembly of complex biomolecular architectures during mussel byssus biofabrication. Nature Communications, 8(1), 14539.

Priemel, T., Palia, R., Babych, M., Thibodeaux, C. J., Bourgault, S., & Harrington, M. J. (2020). Compartmentalized processing of catechols during mussel byssus fabrication determines the destiny of DOPA. Proceedings of the National Academy of Sciences, 117(14), 7613–7621.

Qin, X.-X., Coyne, K. J., & Waite, J. H. (1997). Tough Tendons. Mussel byssus has collagen with silk-like domains. Journal of Biological Chemistry, 272(51), 32623–32627.

Qin, C., Pan, Q., Qi, Q., Fan, M., Sun, J., Li, N., & Liao, Z. (2016). In-depth proteomic analysis of the byssus from marine mussel *Mytilus coruscus*. Journal of Proteomics, 144, 87–98.

Ramsden, C. A., & Riley, P. A. (2014). Tyrosinase : The four oxidation states of the active site and their relevance to enzymatic activation, oxidation and inactivation. Bioorganic & Medicinal Chemistry, 22(8), 2388–2395.

Rzepecki, L. M., Hansen, K. M., & Waite, J. H. (1992). Characterization of a Cystine-Rich Polyphenolic Protein Family from the Blue Mussel *Mytilus edulis* L. The Biological Bulletin, 183(1), 123–137.

Sagert, J., & Waite, J. H. (2009). Hyperunstable matrix proteins in the byssus of *Mytilus galloprovincialis*. Journal of Experimental Biology, 212(14), 2224–2236.

Sagert, J., Sun, C., & waite, J. H. (2006). Chemical subtleties of mussel and polychaete holdfasts. In Biological adhesives (pp. 125–143). Berlin, Heidelberg: Springer Berlin Heidelberg.

Schmitt, C. N. Z., Winter, A., Bertinetti, L., Masic, A., Strauch, P., & Harrington, M. J. (2015). Mechanical homeostasis of a DOPA-enriched biological coating from mussels in response to metal variation. Journal of The Royal Society Interface, 12(110), 20150466.

Silverman, H. G., & Roberto, F. F. (2007). Understanding Marine Mussel Adhesion. Marine Biotechnology, 9(6), 661–681.

Solomon, E. I., Sundaram, U. M., & Machonkin, T. E. (1996). Multicopper Oxidases and Oxygenases. Chemical Reviews, 96(7), 2563–2606.

Stewart, R. J., Weaver, J. C., Morse, D. E., & Waite, J. H. (2004). The tube cement of *Phragmatopoma californica*: a solid foam. Journal of Experimental Biology, 207(26), 4727–4734.

Stewart, R. J., Ransom, T. C., & Hlady, V. (2011). Natural underwater adhesives. Journal of Polymer Science Part B: Polymer Physics, 49(11), 757–771.

Sugumaran, M., Soderhall, K., Iwanaga, S., Vastha, G. (1996). Role of insect cuticle in immunity. In New Directions in Invertebrate Immunology; Eds.; SOS Publications: Fair Haven, NJ, USA, pp. 355–374.

Tamarin, A., Lewis, P., & Askey, J. (1976). The structure and formation of the byssus attachment plaque in Mytilus. Journal of Morphology, 149(2), 199–221.

Teufel, F., Almagro Armenteros, J. J., Johansen, A. R., Gíslason, M. H., Pihl, S. I., Tsirigos, K. D., Winther, O., Brunak, S., Von Heijne, G., & Nielsen, H. (2022). SignalP 6.0 predicts all five types of signal peptides using protein language models. Nature Biotechnology, 40(7), 1023–1025.

Theopold, U., Schmidt, O., Söderhäll, K., & Dushay, M. S. (2004). Coagulation in arthropods : Defence, wound closure and healing. Trends in Immunology, 25(6), 289–294.

Trifinopoulos, J., Nguyen, L.-T., von Haeseler, A., & Minh, B. Q. (2016). W-IQ-TREE : A fast online phylogenetic tool for maximum likelihood analysis. Nucleic Acids Research, 44(W1), W232–W235.

Ullrich, R., & Hofrichter, M. (2007). Enzymatic hydroxylation of aromatic compounds. Cellular and Molecular Life Sciences, 64(3), 271–293.

Van Holde, K. E., & Miller, K. I. (1995). Hemocyanins. In Advances in Protein Chemistry (Vol. 47, p. 1–81). Elsevier.

Vovelle J. (1965). Le tube de *Sabellaria alveolata* (L.) : Annélide polychète Hermellidae et son ciment. Etude écologique, expérimentale, histologique et histochimique. Arch. Zool. Exp. Gén., 106 : 1–187.

Waite, J. H., & Tanzer, M. L. (1981). Polyphenolic Substance of *Mytilus edulis* : Novel Adhesive Containing L-Dopa and Hydroxyproline. Science, 212(4498), 1038–1040.

Waite, J. H. (1985). Catechol Oxidase in the Byssus of the Common Mussel, *Mytilus Edulis* L. Journal of the Marine Biological Association of the United Kingdom, 65(2), 359–371.

Waite, J. H. (1990). The phylogeny and chemical diversity of quinone-tanned glues and varnishes. Comparative Biochemistry and Physiology Part B: Comparative Biochemistry, 97(1), 19–29.

Waite, J. H., Jensen, R. A., & Morse, D. E. (1992). Cement precursor proteins of the reef-building polychaete *Phragmatopoma californica* (Fewkes). Biochemistry, 31(25), 5733–5738.

Waite, J. H., & Qin, X. (2001). Polyphosphoprotein from the Adhesive Pads of *Mytilus edulis*. Biochemistry, 40(9), 2887–2893.

Waite, J. H. (2017). Mussel adhesion – essential footwork. Journal of Experimental Biology, 220(4), 517–530.

Wang, C. S., & Stewart, R. J. (2012). Localization of the bioadhesive precursors of the sandcastle worm, *Phragmatopoma californica* (Fewkes). Journal of Experimental Biology, 215(2), 351–361.

Wang, C. S., & Stewart, R. J. (2013). Multipart Copolyelectrolyte Adhesive of the Sandcastle Worm, *Phragmatopoma californica* (Fewkes) : Catechol Oxidase Catalyzed Curing through Peptidyl-DOPA. Biomacromolecules, 14(5), 1607–1617.

Wang, J., Suhre, M. H., & Scheibel, T. (2019). A mussel polyphenol oxidase-like protein shows thiol-mediated antioxidant activity. European Polymer Journal, 113, 305–312.

Waterhouse R.M., Seppey M., Simão F.A., Manni M., Ioannidis P., Klioutchnikov G., Kriventseva E.V. and Zdobnov E.M. (2018). BUSCO applications from quality assessments to gene prediction and phylogenomics. Molecular Biology and Evolution 35, 543–548.

Yu, J., Wei, W., Danner, E., Ashley, R. K., Israelachvili, J. N., & Waite, J. H. (2011). Mussel protein adhesion depends on interprotein thiol-mediated redox modulation. Nature Chemical Biology, 7(9), 588–590.

Zhao, H., Sun, C., Stewart, R. J., & Waite, J. H. (2005). Cement Proteins of the Tube-building Polychaete *Phragmatopoma californica*. Journal of Biological Chemistry, 280(52), 42938–42944.

