## Supplementary materials for "Abundance, diversity and evolution of tyrosinase enzymes involved in the adhesive systems of mussels and tubeworms"

**This PDF file includes:**

**Supplementary Figs. 1, 3 and 4**

**Other Supplementary Materials for this manuscript include the following:**

**Supplementary Figs. 2.** Detailed maximum likelihood phylogenetic tree (corresponding to main text Fig. 3) showing the distribution of tyrosinase sequences in Lophotrochozoa, based on sequences listed in Suppl. Table 2. The bootstrap values are indicated at each node. The scale bar indicates the substitution rate per site.

**Supplementary Table 1.** List of the tyrosinase protein sequences retrieved from the NCBI database and used for the CLANS analysis and phylogenetic analysis. The reference sequences used in the BLAST analyses are highlighted in bold.

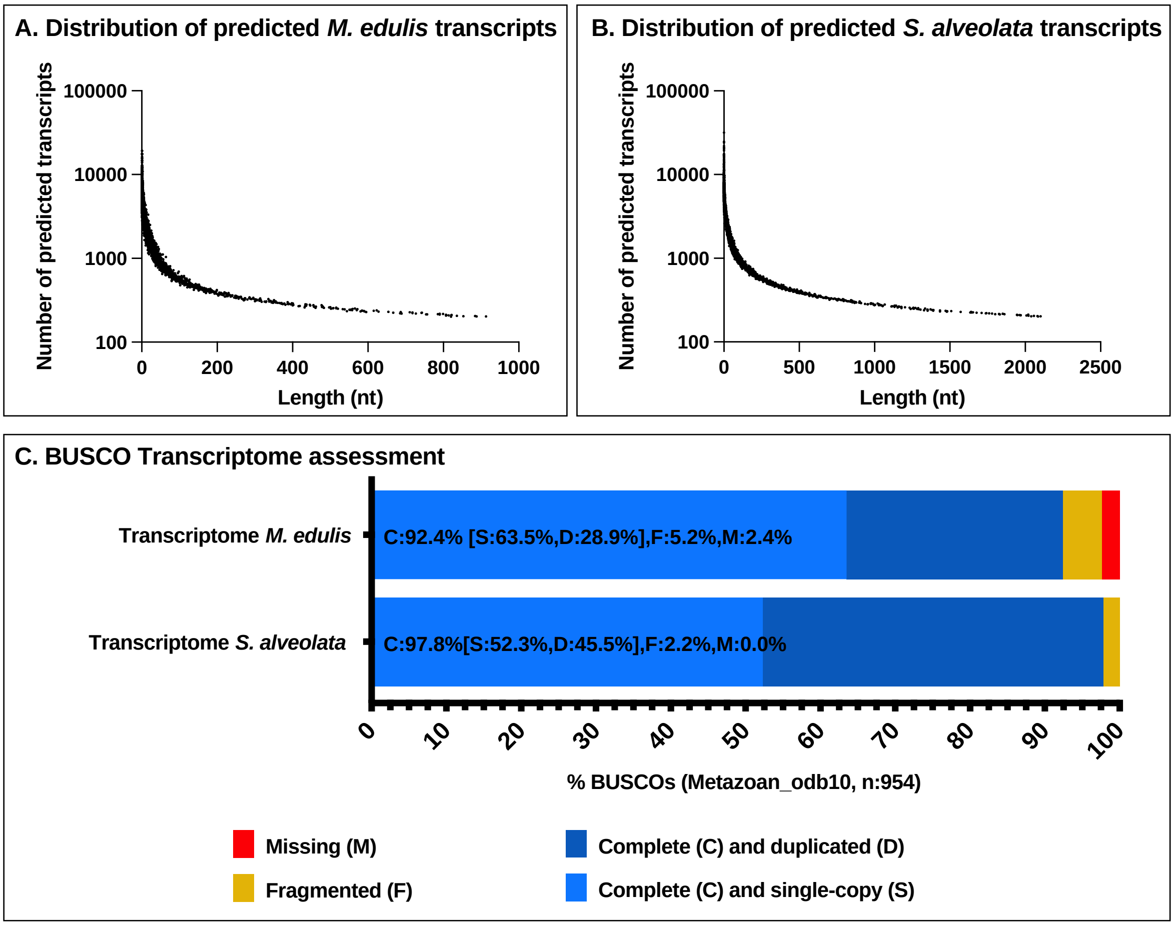


**Supplementary Figure 1. Supplementary Figure 1.** Distributions of predicted transcript length and BUSCO analyses of *S. alveolata* and *M. edulis* transcriptomes.

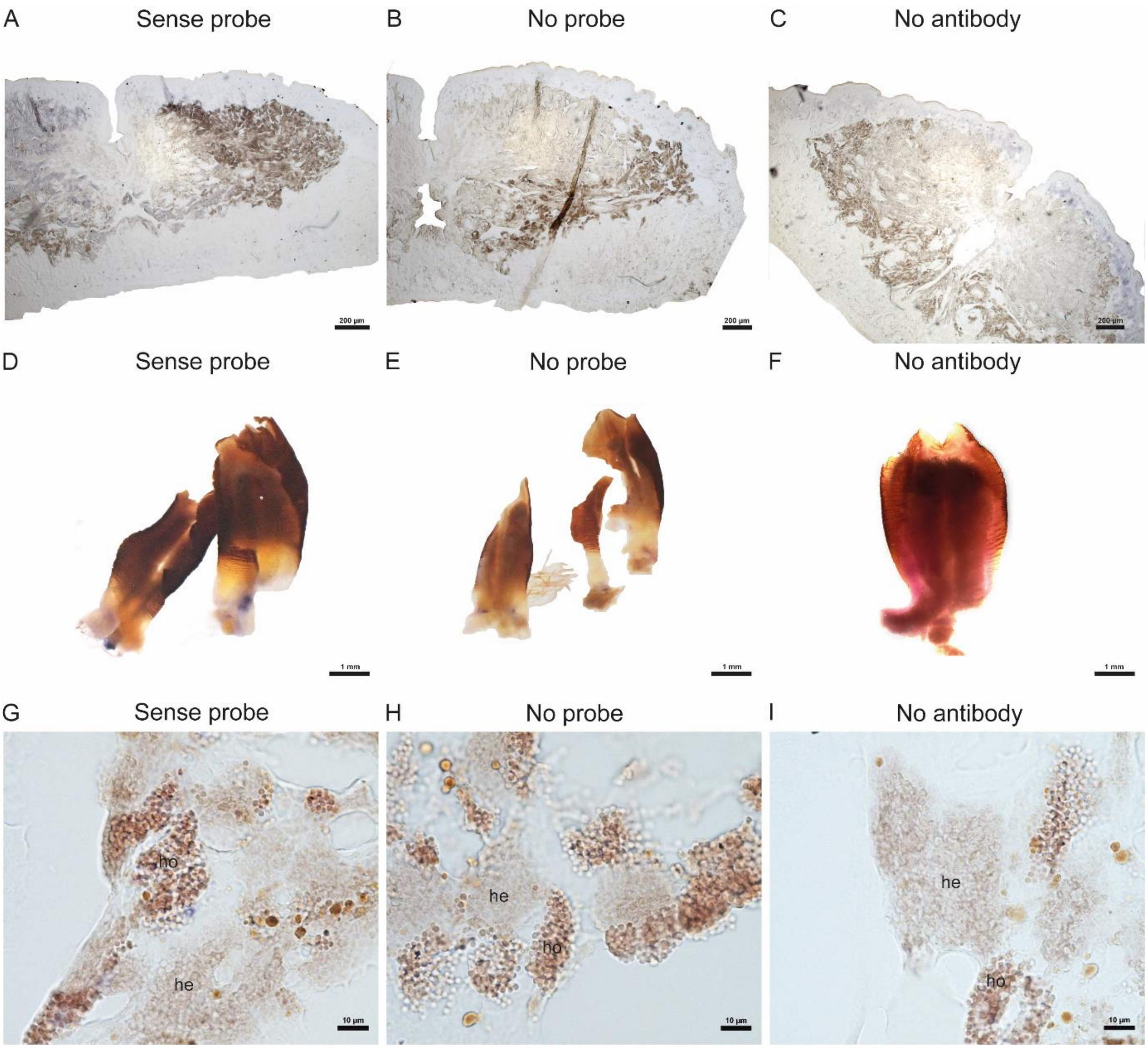


**Supplementary Figure 3.** *In situ* hybridization (ISH) controls with sense RNA probes, without probes and without antibodies. Section (A-C) and whole mount (D-F) ISH on the foot of the mussel *Mytilus edulis*. Section (G-I) ISH parathoracic cement glands of the tubeworm *Sabellaria alveolata*.

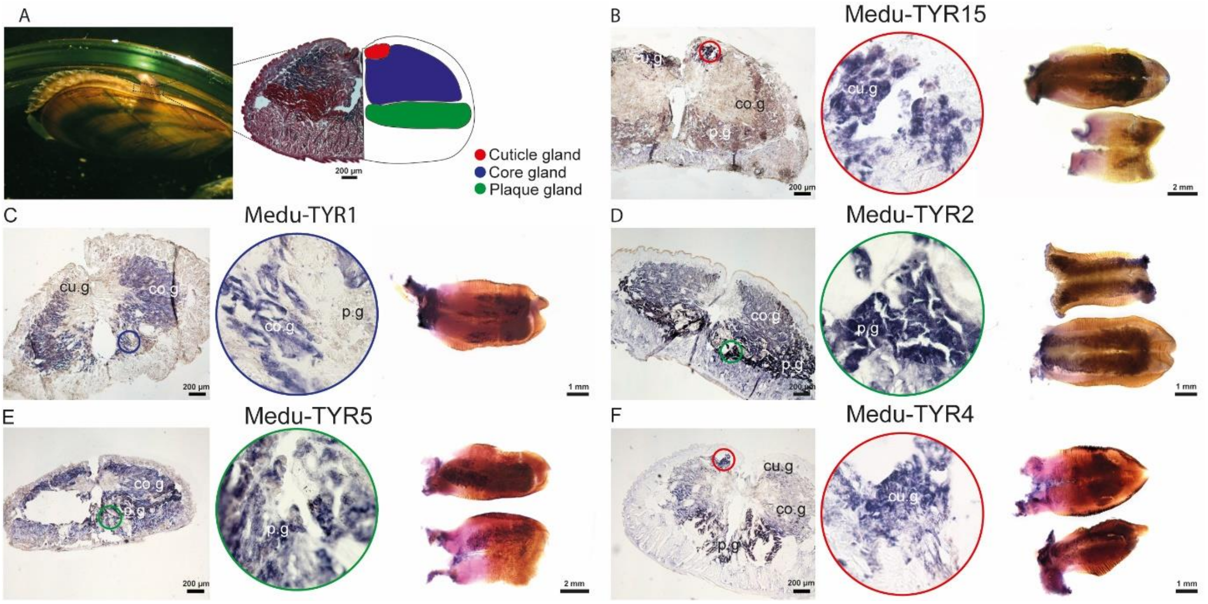


**Supplementary Figure 4.** Expression patterns of selected tyrosinase transcripts in the foot of *Mytilus edulis* (**A**) Illustration showing the mussel's foot and a transverse histological section through this organ stained with Heidenhain's azan stain (left), with a diagram with the arrangement of the different glands in the foot tissues (right). (**B-F**) Section and whole mount *in situ* hybridization of five tyrosinase candidates. Insets in circles show a zoom on the labelled gland. Abbreviations: cu.g – cuticle gland; co.g – core gland; p.g – plaque gland.
