## Supplementary figures and images for "Abundance, diversity and evolution of tyrosinase enzymes involved in the adhesive systems of mussels and tubeworms"

### Suppl Fig 2

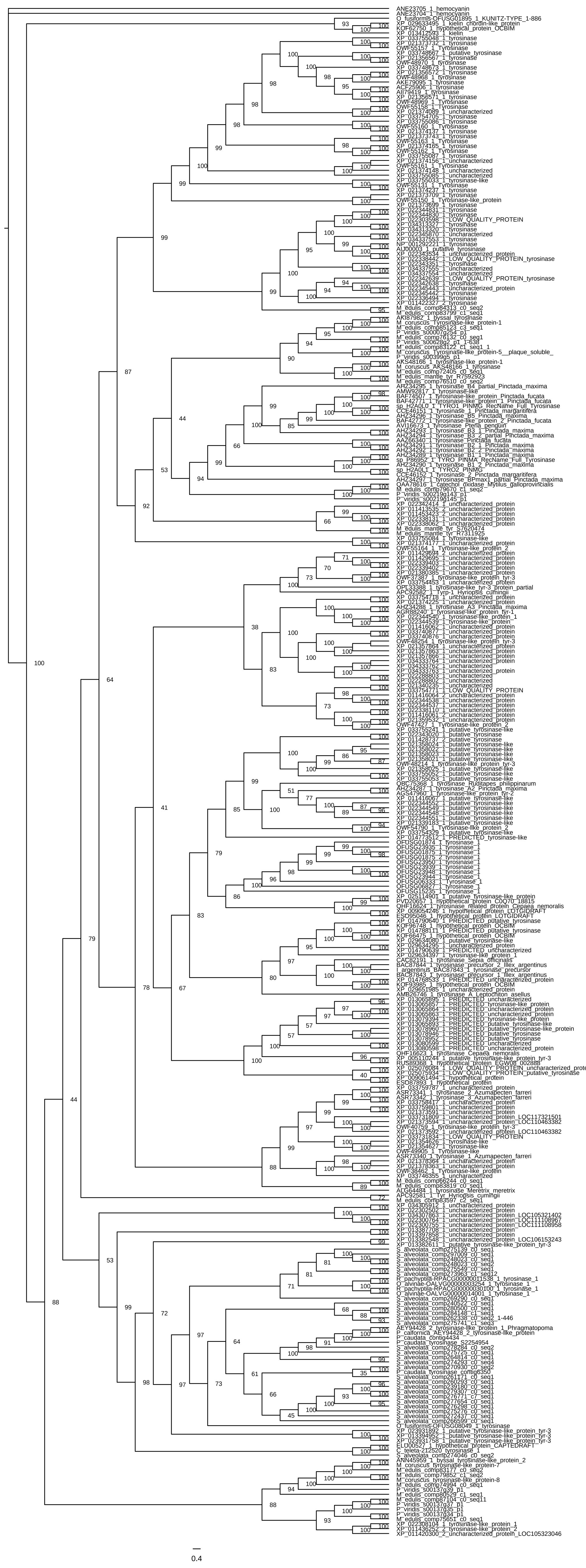
